# A pipeline to create predictive functional networks: application to the tumor progression of hepatocellular carcinoma

**DOI:** 10.1101/605519

**Authors:** Maxime Folschette, Vincent Legagneux, Arnaud Poret, Lokmane Chebouba, Carito Guziolowski, Nathalie Théret

## Abstract

**Background:** Integrating genome-wide gene expression patient profiles with regulatory knowledge is a challenging task because of the inherent heterogeneity, noise and incompleteness of biological data. From the computational side, several solvers for logic programs are able to perform extremely well in decision problems for combinatorial search domains. The challenge then is how to process the biological knowledge in order to feed these solvers to gain insights in a biological study. It requires formalizing the biological knowledge to give a precise interpretation of this information; currently, very few pathway databases offer this possibility.

**Results:** The presented work proposes an automatic pipeline to extract automatically regulatory knowledge from pathway databases and generate novel computational predictions related to the state of expression or activity of biological molecules. We applied it in the context of hepatocellular carcinoma (HCC) progression, and evaluate the precision and the stability of these computational predictions. Our working base is a graph of 3,383 nodes and 13,771 edges extracted from the KEGG database, in which we integrate 209 differentially expressed genes between low and high aggressive HCC across 294 patients. Our computational model predicts the shifts of expression of 146 initially non-observed biological components. Our predictions were validated at 88% using a larger experimental dataset and cross-validation techniques. In particular, we focus on the protein complexes predictions and show for the first time that NFKB1/BCL-3 complexes are activated in aggressive HCC. In spite of the large dimension of the reconstructed models, our analyses over the computational predictions discover a well constrained region where KEGG regulatory knowledge constrains gene expression of several biomolecules. These regions can offer interesting windows to perturb experimentally such complex systems.

**Conclusion:** This new pipeline allows biologists to develop their own predictive models based on a list of genes. It facilitates the identification of new regulatory biomolecules using knowledge graphs and predictive computational methods. Our workflow is implemented in an automatic python pipeline which is publicly available at https://github.com/LokmaneChebouba/key-pipe and contains as testing data all the data used in this paper.

## 1 Background

Hepatocellular carcinoma (HCC) is the most common type of primary liver cancer, which counts for more than 800,000 deaths each year. The incidence of HCC is associated with the development of chronic hepatitis mainly linked to viral infection, alcohol consumption and non-alcoholic fatty liver disease (NAFLD) [1]. Lifestyles [2] and environmental pollution such as particulate matter air pollution [3] also contribute to increase burden in HCC worldwide. HCC is a heterogeneous disease and various genomic alterations associated with the etiologies and the stages of the pathology have been widely documented [4, 5]. A pivotal step in the course of HCC progression is the epithelial-mesenchymal transition (EMT) which allows hepatocytes to transdifferenciate into mesenchymal phenotype whereby escaping to host control and acquiring anti-apoptotic and motility features [6]. Upregulation of EMT markers has been associated with tumor aggressiveness and bad prognosis [7, 8] and associated with inflammatory microenvironment [9]. However, in vivo monitoring of EMT processes remains difficult, due to the spatio-temporal dynamics of these molecular events and the snap-shot nature of biopsies sampling. Understanding EMT to identify new therapeutic targets require integrative and modeling approaches.

To build computational models and integrate experimental data on molecular events, pathway databases can be used. However, despite the fact that numerous publicly available pathway databases currently exist, compiling hundreds of signaling pathways for various biomolecules, very few formal representations linked with automatic inference processes have been proposed so far [10]. The main difficulty appears to be the transfer from the biological representation of a pathway towards a logic knowledge base. Currently, pathway repositories, such as Reactome [11], Pathway Commons [12], KEGG [13], or OmniPath [14] propose their own tools to build graphs. Some of these tools are the Cytoscape [15] plugin CyPath2, PCViz for Pathways Commons; pypath for OmniPath; and ReactomeFIViz [16] for Reactome. However, the resultant graphs are difficult to be transferred into mechanistic models because the notion of causality is often misinterpreted. This misinterpretation is due to the lack of a formal causal representation of biochemical reactions such as protein complexes assemblies. For instance tools such as CyPath2, PCViz, ReactomeFIViz, and pypath model protein complexes using a relation of causality between the protein complex members (protein complex members are the cause and consequence of each other); while in our modeling choice, protein complexes may be triggering other reactions, and their presence is a consequence of the presence of their members. Knowing that signaling cascades are represented by multiple complexes assemblies, this misinterpretation impacts importantly the construction of a mechanistic model when using pathway databases. On the other hand, such tools are very useful to compute topological scores, perform statistical analyses, and to integrate gene expression measurements using enrichment analyses [17]. They remain, however, limited to extract logical consequences of the representation of the biological mechanisms.

The *sign-consistency* framework proposes a way to automatically confront the logic of large-scale interaction networks and genome-wide experimental measurements, provided that a signed oriented network is given and that the experimental measurements are discretized in 3 expression levels (up-regulated, down-regulated and no-change). This framework, introduced in [18], has being applied to model middle- and large-scale regulatory and signaling networks. The two most recent implementations of it are by the means of integer linear programming [19] and logic programming. The latter, implemented in a tool named *Iggy* [20], presents some key aspects: (i) it provides a global analysis applying a local rule which relates a node with its direct predecessors, (ii) it handles a network composed of thousands of components, (iii) it allows the integration of hundreds of measurements, (iv) it performs minimal corrections to restore the logic consistency, and (v) once the consistency is restored, it allows to infer the behaviour (up, down, no-change) of components in the network that were not experimentally measured. In this work we apply this sign-consistency framework to model HCC progression.

Our case study is composed of two input data which were publicly available. First, gene expression data from patients with HCC was extracted from the *International Cancer Genome Consortium* (ICGC) database [21]. Based on the EMT signature from MSigDB [22], HCC samples were clustered into either agressive HCCs (high EMT gene expression) or non-agressive HCCs (low EMT gene expression). Second, the up-stream events of the regulatory events of these genes were obtained by querying automatically KEGG to build a causal model from this database. We used Iggy to study what are the regulatory events that explain the differential expression between low and high aggressiveness from the KEGG interaction knowledge (network of 3,383 nodes and 13,771 edges). We discovered that 146 nodes were predicted, of them 33 refer to gene expression, 110 were protein activities, and 3 were protein complexes activities. 88% of the predictions were in agreement with the ICGC gene expression measurements. Importantly, we predicted the activation of NFKB1/BCL3 and NFKB2/RELB complexes, two critical regulators of NFKB signalling pathway implicated in tumorigenesis. Finally, we proposed a method to discover sensitive network regions that explains HCC progression. This means network components which were highly constrained by multiple experimental data points that could be interesting to target in order to obtain significant changes in the system behavior. We provide a list of 27 nodes discovered by this approach, including TP53.

These results were obtained with a new pipeline developed for this work and freely available online. This pipeline, based on an initial network and a list of genes of interest, allows to extract a functional network based on this list, apply the prediction method described above, and run stability analyses on the result.

## 2 Results

### 2.1 Overview of the pipeline

We introduce key-pipeline: a Python package implementing the workflow for identifying key protein complexes associated to tumor progression. The general pipeline implemented is depicted in Figure 1. It receives as input data: a list of *differentially expressed genes*, a graph describing signed and directed signaling interactions, and a set of *excluded genes* to be filtered out from the graph. Our software allows researchers to: 1) construct the set of *observations*, and the set of *gene names* from a file of *differentially expressed genes* (in CSV format with 3 columns: genes names, log_2_(fold-change) and adjusted *p*-value), see Section 5.1; 2) extract a specific regulatory and signaling network associated with the input genes list from a *signed interaction graph* (based on KEGG regulatory knowledge, see Section 5.2); 3) apply the Iggy tool to compute the *predictions* based on the sign-consistency modeling (see Section 5.3); 4) perform robustness and stability analyses (see Section 5.4); and finally 5) generate plots of these analyses. The pipeline provides a command line interface (CLI), it can be customized by entering file names as arguments. By default, all the steps of the methods will be executed, but the user can run specific steps by using the argument `--steps`. This general pipeline implements all the steps in the workflow described from Section 5.2.2 to Section 5.4.2 and depicted in Figure 1. Each step will output one or more files. In general, the output of one step corresponds to the input of another one. This enables a straightforward application of the workflow for users without programming expertise. We refer the reader to the online documentation for an in-depth description of installation and usage^[1]^.

### 2.2 Integration of Gene Expression in Signaling and Regulatory Network

A first interaction graph was built from the KEGG Pathway database (see Section 5.2). This graph was composed of 41,546 interactions (gene transcriptions, protein signaling, protein formation and complex formations) and 8,861 components (genes, proteins and protein complexes). It is available as input-data with the pipeline^[2]^. Using our pipeline (Figure 1, step 2: Pathrider), the 1913 differentially expressed genes between low and high aggressive tumors were used as input to extract a subgraph from the KEGG pathways graph. Only 209 genes from the 1913 were identified and used to extract upstream regulatory events (Additional file 1: Table S1). In this step the biomolecules associated to the 4,220 genes whose expression is undetectable were filtered out. The resultant graph was composed of 13,771 interactions and 3,383 components (Additional file 1: Figure S1). The content of the graph is available in Additional file 2: File `graph.sif` in SIF format and in Additional file 2: File `graph.cys` as a Cytoscape session.

The final graph contains mostly activations (11,661 versus 2,110 inhibitions); this follows the same activation/inhibition distribution than for the KEGG graph. Only 209 nodes have observations attached to them, provided by the differential analysis of Section 5.1, leaving most nodes unobserved and subject to computational predictions. Finally, the presence of nodes gathering a lot of incoming or outgoing interactions is noteworthy:

- The largest in-degree is 92 (concerning nodes `PIK3R6_prot`, `PIK3CG_prot` and `PIK3R5_prot`);
- The largest out-degree is 79 (concerning nodes `PRKACB_prot`, `PRKACA_prot`);
- Two nodes (`MAPK3_prot` and `MAPK1_prot`) both have the maximal total degree of 107, with 56 incoming and 51 outgoing interactions.

Such “hub” nodes, having an influence to and from a lot of other components, have a high impact on the rest of the network and produce less consensual labelings.

### 2.3 Computational Predictions Validation

We applied the sign-consistency modeling on the final KEGG signed graph obtained in the previous section and the 209 observations derived from the differential analysis (see Section 5.1). Iggy (Figure 1, step 3) returns 146 predictions, that is, couples of (*x*, *s*), where *x* is a node of the graph (either a gene, a protein or a protein complex) and *s* is its consensual sign across all consistent labelings, as given by the *pred*(*x*) function (see Section 5.3). *s* ∈ {−, 0, +} refers to a down-regulation (or inactivation), unchanged behavior, and up-regulation (or activation) of the behavior of biomolecule *x* under the low versus high aggressive tumor comparison. In the case where *x* is a protein complex or a protein, the predicted sign will denote a positive, negative or neutral shift in the effect of the protein or complex towards their targets. In Table 1 we show the 146 obtained predictions after minimal correction, in summary we obtained:

**Table 1.**
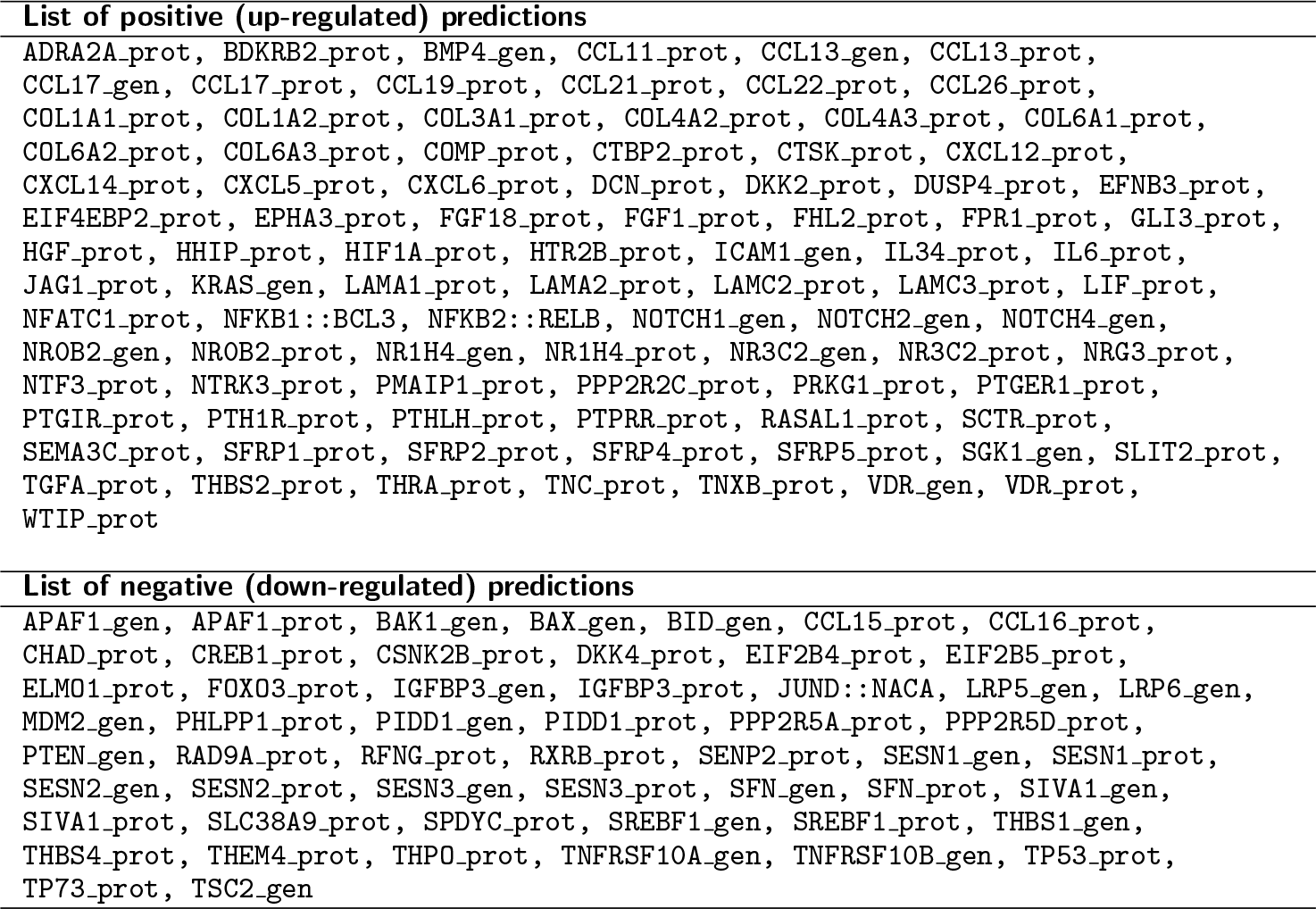
List of all predictions returned by Iggy.

- 92 over-expressions (+): 77 proteins, 13 genes, and 2 protein complexes,
- 54 under-expressions (−): 33 proteins, 20 genes and 1 protein complex.

The list of all predictions is plotted on the KEGG graph in Additional file 1: Figure S2 and on the volcano plot of differential gene expression in Additional file 1: Figure S3. The minimal correction set (MCoS) detected to recover the consistency between the graph causality and the data, was composed of a single repair: adding an influence towards node `PMAIP1_gen` resolves the conflict. This repair is shown in Additional file 1: Figure S4. In the end, 3,026 nodes remain not observed nor predicted. Iggy takes one minute to compute these results on a standard laptop computer^[3]^.

Among the 146 predictions, 143 have a name that matches with a gene name identified in HCC from the ICGC database, but that were not selected in the 1913 genes differentially expressed (< − 0.5 for down-regulated and > 2 for up-regulated in aggressive HCC). If we remove all thresholds and thus consider any positive fold-change as an over-expression, and any negative fold-change as an under-expression, then 82 components predicted + are coherent with the ICGC data and 8 are not; 44 components predicted − are coherent with the ICGC data and 9 are not. This ratio of 88% of matching predictions speaks in favor of our choice of applying Iggy to this specific biological system, with respect to the currently available data in KEGG and ICGC databases. This comparison can be visualized on the volcano plot of gene differential expression in Additional file 1: Figure S3, and is also depicted on the KEGG graph in Additional file 1: Figure S5. The list of predictions not matching with experimental expression data is given in Additional file 1: Table S2.

### 2.4 Impact of data incompleteness on computational predictions

This section presents the results of the two validation analyses applied on the sampling of observations described in Section 5.4. The objective is to observe the impact of data incompleteness in our computational predictions. For this, we observed and tracked across the samples the level of precision and the quality of the information contained in the predictions (See Figure 1, step 4).

#### 2.4.1 Precision Rate

The first approach (see Section 5.4.1) aims at observing the evolution of the predictions precision when using an increasing amount of data-points of the original dataset (see Figure 2). We can observe a clear convergence of the precision score towards 0.88 corresponding to the precision found with the full dataset, which shows that our complete predictions do not lie in a local extremum.

#### 2.4.2 Stability Study

The second approach (see Section 5.4.2) consists in observing “good”, “bad” and “missing” predictions for each of the experiments (samplings < 100%) compared to the 100% sampling. Figure 3 computes the minimum, maximum, median and mean of each such category. Globally, we can observe that the mean and median number of “bad” predictions, that is, predictions that are different with a subset of observations than with the complete set of observations, are really low, below 4% for all samplings. Nevertheless, some samplings show a high proportion of such “bad” predictions. Moreover, the number of “missing” predictions is very high for low samplings, which assesses that there is too little information to obtain complete results. Overall, “bad” predictions tend to decrease after the 65% sampling, along with “missing” predictions that decrease all the way, making “good” predictions mathematically increase.

#### 2.4.3 Insights of the Stability Results

The analyses of the experiments shown in the previous subsections show that the “badly” predicted components for subsets of observations are always the same 28 nodes, listed in Additional file 1: Table S3. These nodes belong to the same region of the graph, which is depicted in Additional file 1: Figure S4. Actually, a group of 27 of these nodes are strongly linked and always change their coloring together. When searching inside the graph topology, one can remark that this group is tightly linked to the node `TP53_prot`, which is also part of the group. This protein acts as a “hub” inside the graph, having a high degree (25 ingoing and 28 outgoing edges). It therefore controls closely, if not directly, a lot of other components that change their sign as soon as it does so, rendering the whole group of predictions unstable. The reason of this instability is that `TP53_prot` directly influences node `PMAIP1_gen` which is involved in the only MCoS repair in our graph: the node `PMAIP1_gen` is indeed observed as over-expressed (+) but 3 other under-expressed (−) observations contradict this one: `CCNG1_gen`, `SHISA5_gen` and `TP73_gen`. This leads to an inconsistency, as explained in Section 2.3. The repair here consists in adding an edge towards `PMAIP1_gen` that models missing information, in order to remove this inconsistency, as shown in Additional file 1: Figure S4. In practise, this renders `PMAIP1_gen` “silent” regarding `TP53_prot`, which then takes the coloring of the other observations (under-expression). Nevertheless, when picking random sets of observations, we sometimes fall in cases where among these 4 observations, only `PMAIP1_gen` is selected; in this case, no repair is needed and `TP53_prot` is predicted as over-expressed, also leading to 26 different predictions in downstream nodes.

Finally, the last unstable node is `PMAIP1_prot`: in the case where `PMAIP1_gen` is part of the randomly picked observations, it is straightforwardly predicted over-expressed while in the converse case, where `PMAIP1_gen` is not part of the observations, it is indirectly influenced by `TP53_prot` and thus predicted under-expressed.

Such unstable predictions can be regarded as not very robust because they are changeable depending on the number of observations taken into account. On the other hand, all other predicted components are stable and can be considered as robust since, when they are predicted, their prediction matches the one obtained using all the observations. The list of stable and unstable predictions is given in Additional file 1: Table S3.

### 2.5 Biological Validation of the Computational Results

Among the computational predictions given in Section 2.3, some of them are of particular interest in regard to the expression data from ICGC. In this section, we detail and validate them biologically.

#### 2.5.1 Activation of NFκB signaling in aggressive HCC

Based on the regulatory model (Additional file 1: Figure S1) and differential expression of mRNA between low and high aggressive HCC (see Section 5.1), the algorithm Iggy predicts the activation of complexes NFKB1∷BCL3 and NFKB2∷RELB and the deactivation of complex JUND∷NACA. By activation of the complexes we mean that in order to explain the high-aggressive versus low-aggressive tumor gene expression datasets, these complexes have to increase their activity. For example, if NFKB1∷BCL3 is activated, then we deduce that its effect on gene IL10 (regulated positively by this complex, Additional file 1: Figure S6) is positive, meaning that the level of gene IL10 may increase if it was only regulated by NFKB1∷BCL3. protein complex prediction is a novel information since it was not present in the initial experimental data of gene expression.

Among them, two complexes are related to NF*κ*B signaling and are predicted as activated: NFKB1∷BCL3 and NFKB2∷RELB. NFKB1, NFKB2 and RELB are three subunits of the transcription factor complex nuclear factor-kappa-B (NF*κ*B) which consist in a homo- or heterodimeric complex formed by Rel-like domain-containing proteins p65 (RelA), RelB, c-Rel, p50 (NFKB1), and p52 (NFKB2). The NF*κ*B signaling system acts through canonical and non canonical pathways which are induced by different extracellular signals [23]. The canonical pathway can be induced by TNF-*α*, IL-1 or LPS stimulation and requires NF-kappa-B essential modulator (NEMO) while the non-canonical pathway is induced by other ligands such as CD40 ligand (CD40L), receptor activator of nuclear factor kappa-B ligand (RANKL), B-cell activating factor (BAFF) and lymphotoxin beta (LTb). Upon ligand binding to its receptor, the signaling cascades control the degradation of IkB proteins (inhibitor of NF*κ*B) and precursor processing including NFKB1 (p105) and NFKB2 (p100) which are proteolytically activated to p50 and p52 respectively. B-cell chronic lymphatic leukemia protein 3 (Bcl3) is a member of IkB family that are inhibitors of NF*κ*B members. BCL3 associates with NF-kappa B in the cytoplasm and prevents nuclear translocation of the NFKB1 (p50) subunit. When phosphorylated, BLC3 is activated and associates with NFKB1 in the nucleus to regulate NF*κ*B target genes [24]. NF*κ*B system is involved in the regulation of numerous biological processes including inflammation, cell survival and development. Regarded as protective against aggression from environment in normal physiology, alteration of NF*κ*B signaling pathways has been associated with various diseases such as inflammatory disease and cancer [25, 26]. In HCC, NF*κ*B pathway was shown to be deregulated in tumor and underlying fibrotic livers [27, 28]. Notably, increased expression of p50 and BCL3 has been reported in tumors compared with adjacent tissues [29] and p50 expression was associated with early recurrence of HCC [28].

In order to evaluate our predictions about the activation of NFKB1∷BCL3 and NFKB2∷RELB complexes, we thought to search for expression of genes regulated by these complexes. For that purpose, we take advantage of the NF*κ*B-dependent signature available in MSigDB [22, 30]. We selected the `HALLMARK_TNFA_SIGNALING_VIA_NFKB`^[4]^ signature which contains 200 genes regulated by NF*κ*B in response to TNF. As shown in Additional file 1: Figure S7A, we demonstrated that these genes were more expressed in high aggressive HCC when compared with low aggressive ones supporting the activation of NF*κ*B signaling. More specifically, we searched for expression of genes targeted by NF*κ*B-non-canonical pathway, including the cytokines CCL19 and CCL21. These genes are regulated through the activation of NFKB2∷RELB complexes and their expression was increase in high aggressive HCC thereby confirming the prediction (Additional file 1: Figure S7B).

Another prediction was the down-regulation of JUND∷NACA complex that was previously demonstrated to regulate osteocalcin [31]. This prediction is mainly conditioned by osteocalcin (BGLAP) expression data that was found down-regulated in the aggressive HCC (−1.3 fold-change between aggressive versus non-aggressive HCC). Such observations are in accordance with previous reports showing that osteocalcin was down-regulated in the serum of HCC patients when compared with healthy controls [32]. As shown in Additional file 1: Figure S8A, we showed that both JUND and NACA gene expressions were down-regulated in aggressive HCC supporting the prediction of down-regulation of the complexes JUND∷NACA. Importantly, the targets of JUND∷NACA complex including LRP5 and LRP6 genes were predicted as down-regulated by our model (Additional file 1: Figure S6). The down-regulation of LRP5 in aggressive HCC was validated in HCC data but was not significant for LRP6 probably due to the low level of gene expression (Additional file 1: Figure S8B). According with this, the up-regulation of LRP6 through JUND∷NACA complexes was clearly demonstrated in osteoblasts [33].

To conclude, model predictions were validated by data analyses and are in accordance with the literature. Importantly, this is the first report describing the activation of NFKB2∷RELB complex and the down-regulation of JUND∷PACA complex in aggressive HCC.

## 3 Discussion

The understanding of tumor progression dynamics is extremely difficult when considering the snap-shot nature of data from patients. However, compiling information from a wide spectrum of tissue samples can be used for modeling evolutive stories. The complexity of molecular events implicated in hepatocellular carcinoma progression is directly associated with its various etiologies that differently contribute to tumor initiation, growth and evasion. During last decades, multiscale omics data analysis of genome and proteome allowed to explore molecular networks associated with HCC and mathematical models have been developed namely to predict cancer cell behavior [34]. Accordingly, an elegant discrete model was developed by [35] to explore TGF-β signaling pathway during epithelio-mesenchymal transition in HCC. However, HCC results from complex interactions between the tumor cells and the microenvironment involving stromal cells and extracellular matrix. Molecular biological data from tumor tissues recapitulate all this information and we need to build an unique large-scale model without *a priori* to take into account such complexity. For that purpose, we propose here an original approach aiming at integrating experimental data on a regulatory graph extracted from the KEGG database to predict new markers and regulators of HCC progression.

Based on EMT gene expression signature from MSigDB [22] we first separated low from high aggressive HCC samples stored in the ICGC database [21] and next we sought to predict the regulatory pathways implicated in this transition. For that purpose, we built a model by querying the KEGG database using the KEGG API to extract an initial network. We have implemented a tool, Pathrider, to allow us extracting a directed and signed sub-network, from the previously obtained network, by using the up-stream events of a list of *target genes*. Importantly, our modeling choices allowed us to connect protein complexes to their members, and to label network nodes of type *gene* and *protein*. This separation of concepts is particularly valuable when modeling gene expression.

The publicly available knowledge base KEGG, gathering curated signaling and regulatory processes, is well structured to automatically extract mechanistic models from it. In particular: (i) the information concerning gene transcription and signaling modifications is differentiated, (ii) the network nodes identifiers are unique, and (iii) the biological processes, such as phophorylation or gene-regulation, are clearly represented.

Using Iggy, it was possible to confront the logic of a large-scale KEGG network (3,383 nodes, 13,771 edges) to the expression of genes differentially expressed between high-aggressive and low-aggressive HCC. In this context, we were able to propose an integrated model of HCC progression and to predict the regulation of new biomolecules including genes, proteins and complexes. A major finding is that the model predicted the behavior of 146 network components that were associated with the progression of tumors. 88% of the computation model predictions were validated with the ICGC data-set and by using cross-validation techniques, thereby demonstrating the quality of the model. Conversely, 12% of the predictions did not match the experimental data, however 10 of these components are part of gene/protein couples leading to linked predictions. In addition, all of these components but one had a low expression change (less than 1 in absolute value) along with a high *p*-value (above 10^−2^) that might explain the inconsistency. The remaining one is `THBS1_gen` (thrombospondin 1 gene) with a fold-change of 1.996, and is also part of the cluster of unstable predictions depicted in Section 2.4.3. Indeed, we discovered a subset of 28 network nodes that were very sensitive to the experimental data. That is, they were strongly constrained by a subset of experimental observations. We notice that these nodes behave as hubs in the network, and can be candidate to experimental stimulation or inhibition in order to affect the system behavior.

The most interesting prediction was the activation of protein complexes related to NF*κ*B signaling since complexes formation is directly responsible for signal transduction [36]. While the role of NF*κ*B signaling pathway has been widely documented in chronic liver disease [37], the activation of NFKB1/BCL-3 complexes in aggressive HCC has never been reported. The I*κ*B protein BCL-3 acts both as a co-activator that form complexes with NFKB1(p50) dimers to promote genes [38] and as a co-repressor of gene transcription by stabilizing P50 homodimers on DNA promoters [39]. Predicted activation of such complexes in aggressive HCC revealed the ambivalent role of NFKB-mediated inflammatory response during the course of tumor progression [40].

## 4 Conclusion

The present study is general to be applied to other biological data from cancers or other disease. In the future, we would like to use logic programming to target the combinatorics of sensitive regions in a regulatory graph with respect to gene expression profiles, in order to propose regulatory elements for clinical therapy. Another perspective is to apply our method to subsets of patients, and observe if there are clusters of patients that have specific computational model signatures for HCC progression.

## 5 Methods

### 5.1 Identification of gene differentially expressed between low and high aggressive HCC

Normalized HTseq counts and clinical data were retrieved from the LIHC-US project^[5]^ (NCI, TCGA-LIHC). These files were downloaded on 2016-07-19, corresponding to release 21. At this date, LIHC-US dataset comprised 294 donors and 345 samples; among them, we selected samples corresponding to solid primary tumors, based on clinical data, by selecting entries containing the expression “`Primary tumour - solid tissue`” in the specimen table (7^th^ field). This allowed selecting one sample for each of the 294 donors. Data retrieval and filtering workflow is detailed in Additional file 3: File `dataset_filtering.sh`.

From this filtered dataset, we extracted a two-dimensional table of expression values (converted in log_2_) for 20,502 genes in 294 LIHC samples. Based on the bimodal distribution of these expression values, we discarded genes whose expression is undetectable (4,220 genes), keeping 16,282 genes. Expression values were normalized by the median value in each sample. Based on the established link between epithelialmesenchymal transition (EMT) and tumor aggressiveness [41], we used the MSigDB [42] set of 200 genes termed `HALLMARK_EPITHELIAL_MESENCHYMAL_TRANSITION`^[6]^ from the Broad Institute as a molecular signature of aggressiveness. From the LIHC dataset, we extracted a table of expression values for 195 entries of this EMT signature for each of the samples (5 genes were undetectable). Based on the expression values of the EMT signature, LIHC samples were classified (hierarchical clustering of euclidean distances) into three groups termed *low_EMT* (70 samples), *medium_EMT* (154 samples), and *high_EMT* (70 samples). The result of this clustering analysis is available in Additional file 1: Figure S9. Samples corresponding to the *medium_EMT* group were discarded and a differential expression analysis was performed by computing a non-parametric Mann-Whitney test for all the 16,282 genes between the *low_EMT* and *high_EMT* groups. *p*-values were adjusted for multiple analyses by the Benjamini & Hochberg method. The volcano plot of Additional file 1: Figure S10 represents fold-changes (log_2_) against adjusted *p*-values (− log_10_) and the raw data are available online as input data of the pipeline^[7]^.

We focused on genes with an adjusted *p*-value below 10^−5^. Genes with a log_2_(fold-change) greater than 2 were considered as over-expressed (821 genes), whereas those with a log_2_(fold-change) lower than 0.5 were considered as underexpressed (1,092 genes). Together, these 1913 differentially expressed genes, listed in Additional file 2: File `diffexp_filtered.csv` were subsequently used to extract a regulatory network, as explained in Section 5.2, and then used as observations for the coloring propagation process, as detailed in Section 5.3. The full list of differentially expressed genes is available in Additional file 2: File `diffexp_full.csv`. The workflow of data clustering and differential analysis is available in Additional file 3: File `diffexp_and_clustering.R`.

To further validate the clinical relevance of the groups of HCC samples identified by the hierarchical clustering method, we compared this classification obtained with the EMT signature to a classification obtained with markers established as predictive of recurrence, which is a major clinical outcome of tumor aggressiveness. For that purpose, we used the Seoul National University recurrence (SNUR) signature [43] that previously permitted to classify HCC samples from TCGA database [44] and we compared clusters identified by hierarchical clustering method with SNUR groups. Note that LIHC primary tumors correspond to 294 samples but only 183 are annotated with SNUR score. Clustering methods were applied to all the 294 samples used in this study and comparisons of clustering classes were made for the 183 samples. When hierarchical groups were compared to SNUR groups, we found a *χ*^2^ test *p*-value of 3.81 × 10^−14^, with 9% of class 1 (low EMT) belonging to the low-recurrence group, and 83% of class 3 (high EMT) belonging to the high-recurrence group. Together, these data demonstrate the accuracy of clustering method to identified low and high aggressive HCC samples.

### 5.2 Building a signed interaction graph from the KEGG Pathway database

For this case study, we used a human signaling network derived from the KEGG Pathway database [13]. 154 human signaling pathways were fetched using the KEGG API and converted to SIF (Simple Interaction Format) in order to provide KEGG’s knowledge as a graph representation. This section summarizes how this network was built. A more in-depth description is available in Additional file 1 as a Supplementary Material & Methods. This step of automatic reconstruction of a causal graph from KEGG constitutes one of the novel contributions of the methodological results of this paper.

#### 5.2.1 Signed interaction graph built from KEGG’s regulatory knowledge

To model the KEGG regulatory knowledge we imposed a distinction between nodes representing genes and nodes representing proteins. In the KEGG Pathway database, this distinction is implicitly embedded in the relation types, particularly `PPrel` edges (protein-protein relations) and `GErel` edges (gene expression relations).

`PPrel` edges indicate that both source and target nodes are proteins. `GErel` edges indicate that source nodes are transcription factors and that target nodes are genes. Therefore, to explicitly differentiate genes and proteins, the source nodes of `GErel` edges were suffixed with `_prot` and the target nodes were suffixed with `_gen`. Concerning `PPrel` edges, both the source and target nodes were suffixed with prot. Differently to what proposed by KEGG maps, we modeled protein complexes explicitly by imposing two relations:

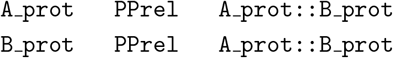

where `A_prot∷B_prot` refers to the protein complex formed by proteins `A` and `B`.

Furthermore, in order to link genes and their products, a relation type (initially absent in the KEGG KGML model) was added: the `GPrel` type (gene-protein relations). For each node *C* modeling a protein, a `GPrel` edge starting from its corresponding gene and ending on *C* was added:

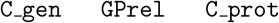

These added nodes therefore model the corresponding gene products and the `GPrel` edges model the protein formation given a gene expression, as illustrated in Additional file 1: Figure S11 and Figure S12. Note that without `GPrel` nodes, the graph is much more disconnected and the predictions are fewer and of worse quality, as showed in Additional file 1: Table S4 and Table S5.

In addition to their relation types, the edges are annotated in KEGG with keywords bringing details about the modeled interactions. Therefore, edge signs (role of activator or inhibitor) were inferred using these keywords.

Altogether, the human signaling network extracted from KEGG was represented as a signed interaction graph composed of protein signaling interactions, complex formations, gene expression regulations and gene-protein relations; accordingly, the nodes of this graph refer to genes, proteins and protein complexes. This decomposed representation allowed us to map on this network data which corresponded only to gene expressions, without assuming that gene expression correlates to protein activity. This KEGG signed and directed graph is available as input data of the pipeline implemented in this work^[8]^.

#### 5.2.2 Extracting up-stream signaling pathways

For this work, we implemented *Pathrider* ^[9]^ in order to extract a subgraph from the KEGG generic human signaling network (obtained in the previous section). Given a list of genes and a network, Pathrider will keep only the signaling pathways of the network regulating the list of genes, that is, the upstream paths of the nodes in the network that represent these genes. Pathrider will also filter out the biomolecules in the graph (genes, proteins or protein complexes) that appear in a *list of excluded genes*, which in practice refer to genes whose expression is undetectable. This tool is included in the automatic pipeline we propose in this work (see Section 2.1).

### 5.3 Sign consistency - Iggy

Let *G*(*V*, *E*, *σ*) be an interaction graph, where *V* represents the set of nodes, *E* the set of edges, and *σ*: *E* → {+, −} a labeling (activation or inhibition) of the edges; and given experimental observations (e.g. gene expression profile) (*S*, *μ*), defined by the set of experimentally measured biomolecules *S* ⊂ *V* and the mapping *μ*: *V* → {−, 0, +}. The sign-consistency principle, implemented in Iggy [20], defines the rules to integrate interaction and experimental knowledge. In order to do this, we look for total labelings *μ*^*t*^: *V* → {−, 0, +} that satisfy the following sign-consistency constraints (see Figure 4):

- The observations must keep their initial labelings.
- Each labeling + or − must be justified by at least one predecessor.
- Each labeling 0 must have only predecessors labeled as 0 or a couple of + and − labeled predecessors.

Given a particular biological instance for *G* and (*S*, *μ*), it usually happens that many total labelings *μ*^*t*^, satisfying the constraints, are proposed. For a set *V* of nodes in our network and a set *M* of total labelings consistent with our observations, we define the prediction function *pred*: *V* → {+, −, 0, ∅} as follows:

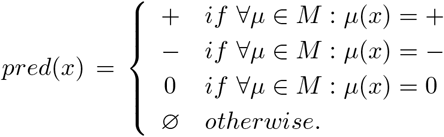

This prediction function is an output of the sign-consistency modeling approach, and it can be seen as an inference mechanism that generates a sign for a node *x* only if in all the consistent total labelings it was assigned the same sign among {−, 0, +}. There may be nodes in *V* with non prediction value (∅) thus meaning that no certain conclusion was possible to be inferred for them. Biologically, this prediction function will allow us to label non experimentally observed nodes, meaning that their shift of expression or activity can be inferred thanks to their connectivity to other observed biomolecules in the graph.

Another possible output of the sign-consistency approach is a list of conflicts, in the case where (*S*, *μ*) is not compatible with *G*. It particularly signals a conflict between the sign of some biomolecules and the interaction network. One way to fix such conflicts is to add artificial interactions in the network. Iggy allows to automatically add a minimal number of such repairs, called *minimal correction set* (MCoS). If several possibilities of repairs are possible, Iggy will compute them all and the final set of predictions will correspond to the union of the predictions obtained after each possible repair.

Given the combinatorial nature of this analysis, Iggy is implemented in Answer Set Programming, in particular using grounder Gringo 3.0.5 and solver Clasp 3.1.3. For this work, the reasoning of Iggy, parametrized with the mentioned constraints and the MCoS repairs, is added to the automatic pipeline we provide.

#### 5.3.1 Modeling our case-study using sign-consistency: inputs and outputs of Iggy Inputs

The signed interaction graph *G*(*V*, *E*, *σ*) was obtained from the KEGG Pathways database as explained in Section 5.2. The gene expression profile (see Section 5.1) is composed of experimental knowledge of 821 over- (sign ‘+’) and 1092 under-expressed (sign ‘−’) genes in in high aggressive tumors compared to low aggressive tumors. From these differentially expressed genes, only 209 were found in the KEGG graph matching nodes with a suffix ‘ gene’. Thus, the set of observations (*S*, *μ*) used for the sign-consistency analysis was composed of 209 elements.

##### Outputs

Following the sign consistency analysis, Iggy proposed predictions under minimal MCoS repairs. The complete results are discussed in Section 2. It is important to recall that since the graph *G*, obtained from KEGG, is composed of nodes which represent genes, proteins, and protein complexes. The prediction function *pred*(*x*) computed by Iggy for *x* ∈ *V* will assign signs mainly to protein and protein complex nodes. In this way Iggy will allow us to infer the activity or expression shifts of unmeasured biomolecules of the system.

### 5.4 Computational Validation of the results

Recall that to create the over- and under-expressed genes between low and high aggressive tumors (see Section 5.1) we used thresholds of +2 and −0.5 on the value of log_2_(fold-change). In this section, we aim at checking if these thresholds are justified. To do this, we computed “sub-predictions”, that is, predictions on the same extracted graph of Section 5.2 but with subsets of observations. To generate these subsets of observations, we considered a range of samplings, from 10% to 95% of the complete observation set, with a step of 5%. For each sampling of *x*%, 100 experiments were conducted, where an experiment consisted in randomly picking *x*% of the over-expressed observations (+) and *x*% of the under-expressed observations (−), and computing the predictions on this subset of observations. The results are 1,800 such subsets of observations, and as many computed sets of predictions on the nodes of the graph, hereafter called *sub-predictions*. These sub-predictions have been exploited in two ways:

1. by comparing said sub-predictions with the available gene expression data from ICGC that were already used for the differential analysis (Section 5.4.1), and
2. by comparing said sub-predictions with the final predictions obtained with 100% of observations to witness their variability (Section 5.4.2).

Both approaches are explained below; they are implemented and added to the automatic pipeline we propose in this work.

#### 5.4.1 Recovery rate of the sub-predictions

We computed a normalized score by counting the number of predictions matching the related experimental fold-change from the ICGC data. For each experiment result, this score *s* is given by the formula: *s* = *m*/*t* where *m* is the number of matching predictions, that is, positive predictions with positive fold-changes and negative predictions with negative fold-changes, and *t* is the total number of predictions. This allows us to assess the ability of our model to recover from missing information (here, observations).

#### 5.4.2 Stability of the sub-predictions

In order to look at the stability of the predictions made on subsets of observations, we also compared them to the final predictions using 100% of the observations. For each predicted node in the 100% sampling set, and for each of its corresponding sub-prediction in a lower sampling set:

- If the node is predicted and the prediction matches the one at 100% sampling, this is considered a “good” prediction.
- If the node is predicted but the prediction is not the same as for 100% sampling, this is considered a “bad” prediction, thus representing mathematical non-monotonicity and biological sensitive components or potential targets.
- If the node is not predicted, this is called a “missing” prediction.

Counting the elements and observing the evolution of these categories allows us to witness if lower samplings converge to the final sampling or not, independently of any exterior data such as expression data.

## Supporting information

Additional tables, figures and explanations

Input data and results of the application of our pipeline regarding hepatocellular carcinoma progression

Additional information to generate differentially expressed genes from ICGC database

## List of abbreviations

HCC: Hepatocellular carcinoma
ICGC: the International Cancer Genome Consortium
NAFLD: non-alcoholic fatty liver disease
EMT: epithelial-mesenchymal transition
CLI: command line interface
MCoS: minimal correction set.

## DECLARATIONS

### Ethics approval and consent to participate

Not applicable.

### Consent for publication

All the authors are aware of, and agree to, the contents of the manuscript.

### Availability of data and materials

The authors confirm that the data supporting the findings of this study are available within the article and its supplementary materials.

### Author’s contributions

MF implemented the different parts of the pipeline presented in this work, except for Iggy and Pathrider. VL produced the differential expression data and participated in the biological validation from litterature. AP produced the KEGG graph extraction and the Pathrider/Stream tool. LC wrote the main script automating the calls to the different parts of the pipeline. CG co-lead the project and gave expertise on the computer science part of this work. NT co-lead the project and participated in the biological validation from litterature. All authors have equally contributed to the writing of the manuscript.

## Acknowledgements

The authors wish to thank Anne Siegel for her fruitful discussions and comments.

---

**Figure 1 Schema describing the pipeline for building networks and predicting regulatory nodes** (1) Using a list of differentially expressed genes, construct the set of gene names and the corresponding set of observations (a sign is attributed for each gene: + when fold change > 2 or − when fold change < 0.5 and adjusted p-value < 10^−5^); (2) Extract the upstream/downstream signaling pathways for the set of genes from the *signed interaction graph* using *Pathrider*, a tool developed in our team to this purpose. Given a list of *excluded genes* (such as invariant genes), *Pathrider* filters these genes to reduce the graph size; (3) Check the sign consistency of our datasets to produce signed predictions for unmeasured biomolecules using *iggy* tool; (4) Validate the *predictions* made by *iggy* by computing sub-predictions (prediction 1, 2…n) using a sub-set of observations (by default, it starts sampling from 10% to 95% of observations with a step of 5% and a number of execution equal to 100), then compare it firstly with the *differentially expressed genes*, and in a second time with the *predictions* obtained with all the set of *observations*; and (5) Plot the precision scores for each sub-sets of the *observations*, and the stability of the *prediction* compared to the predictions of the entire set of *observations*.

**Figure 2 Precision scores of predictions obtained on samplings of the observations** Boxplots of the precision scores (ordinate) of the predictions obtained with 100 randomly picked samplings (abscissa) of observations. Each box plot at abscissa *x* represents the distribution of the precision scores of the predictions obtained when using only *x*% of the observations. The point at 100% represents the prediction score of the predictions when using the complete set of observations.

**Figure 3 Stability of the predictions for subsets of observations** This figure summarizes the stability of the predictions for all samplings of the observations, compared to the final predictions with all 100% of observations. “Good” predictions (matching the 100% predictions) are depicted in green, “Bad” predictions (predicted differently than the 100% predictions) in red and “Missing” predictions (not predicted) in blue. For each category, four curves are plotted representing, from top to bottom, the maximum, median, mean and minimum number of predictions of this type. Curves are normalized to the number of predictions obtained for each set of sampled data.

**Figure 4 Consistency constraints**. Given a signed graph, where green edges depict activations, and red edges, inhibitions, two partial labelings for the nodes *A* and *B* are proposed in the first and second rows. In both cases there are 3_2_ different possible labelings for C and D taking 3 signs {+,−, 0} corresponding to colors green, red and blue, respectively. We depicted here only one consistent and one inconsistent scenario according to the sign-consistency constraints.

## Additional Files

### Additional file 1 — Additional tables, figures and explanations

This PDF file contains additional figures and tables related to all parts of this manuscript, along with a detailed explanation of the KEGG graph extraction that was summarized in Section 5.2.

### Additional file 2 — Input data and results of the application of our pipeline regarding hepatocellular carcinoma progression

This archive contains input and output data related to hepatocellular carcinoma agressiveness that were used in this work to illustrate the benefits of our pipeline. The input data consists of differentially expressed genes (in CSV format) and the KEGG graph extraction (in SIF format). The output data consists in a Cytoscape session to explore the graph and the computational prediction results along with dynamic plots of the results (volcano plots, precision and stability studies, in HTML format).

### Additional file 3 — Additional information to generate differentially expressed genes from ICGC database

This archive contains a SH script to filter dataset and a R script for data clustering and differential analysis

[1] File README.md at https://github.com/LokmaneChebouba/key-pipe/

[2] Input data are available at https://github.com/LokmaneChebouba/key-pipe/tree/master/example

[3] Laptop computer containing an Intel Core i7-5600U CPU with 4 threads of 2.60GHz and running Fedora 27 64 bits.

[4] Id: M5890, available at http://software.broadinstitute.org/gsea/msigdb/geneset_page.jsp?geneSetName=HALLMARK_TNFA_SIGNALING_VIA_NFKB

[5] All ICGC data used in this work are publicly available at https://dcc.icgc.org/releases/release_21/Projects/LIHC-US

[6] Id: M5930, available at http://software.broadinstitute.org/gsea/msigdb/geneset_page.jsp?geneSetName=HALLMARK_EPITHELIAL_MESENCHYMAL_TRANSITION

[7] Files `GSEA_EMThigh_vs_EMTlow_diffexp.csv` (differential expression) and `LIHC_primary_weakly-expressed_genes.txt` (blacklist of weakly expressed genes) at https://github.com/LokmaneChebouba/key-pipe/tree/master/example.

[8] File hsa-2345-symb-nomulti-split-func-sign.sif at https://github.com/LokmaneChebouba/key-pipe/tree/master/example.

[9] Available at: https://github.com/arnaudporet/pathrider

[10] https://www.genouest.org

