## Additional tables, figures and explanations for "A pipeline to create predictive functional networks: application to the tumor progression of hepatocellular carcinoma"

This file contains supplementary Material & Methods, Supplementary Figures and Supplementary Tables related to the article entitled *A pipeline to create predictive functional networks: application to the tumor progression of hepatocellular carcinoma.*

#### Supplementary Methods

**Building the Signaling Network from the KEGG Pathway Database 15**

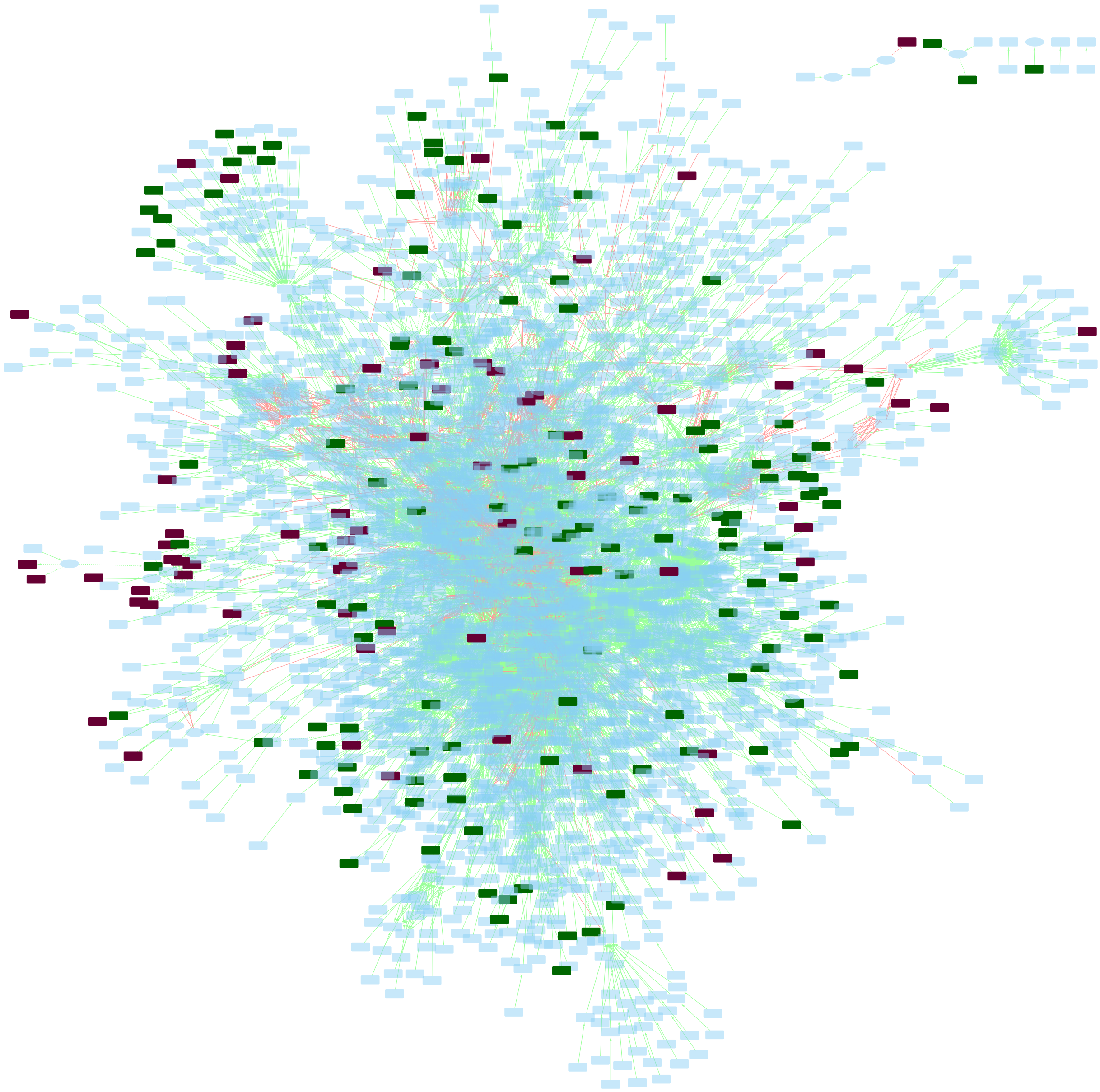

Figure S1: Graph extraction of KEGG by considering only the predecessors of the over- and under-expressed genes of Section 5.1, as explained in Section 5.2. The dark green nodes are observed as over-expressed and the dark red nodes are observed as under-expressed. Green edges are activations and red edges are inhibitions. Plain lines model signaling while dashed lines model regulation interactions. This graph is available with labeled nodes as a Cytoscape session in Additional file 2: `graph.cys`, with style `1-Graph-extraction`.

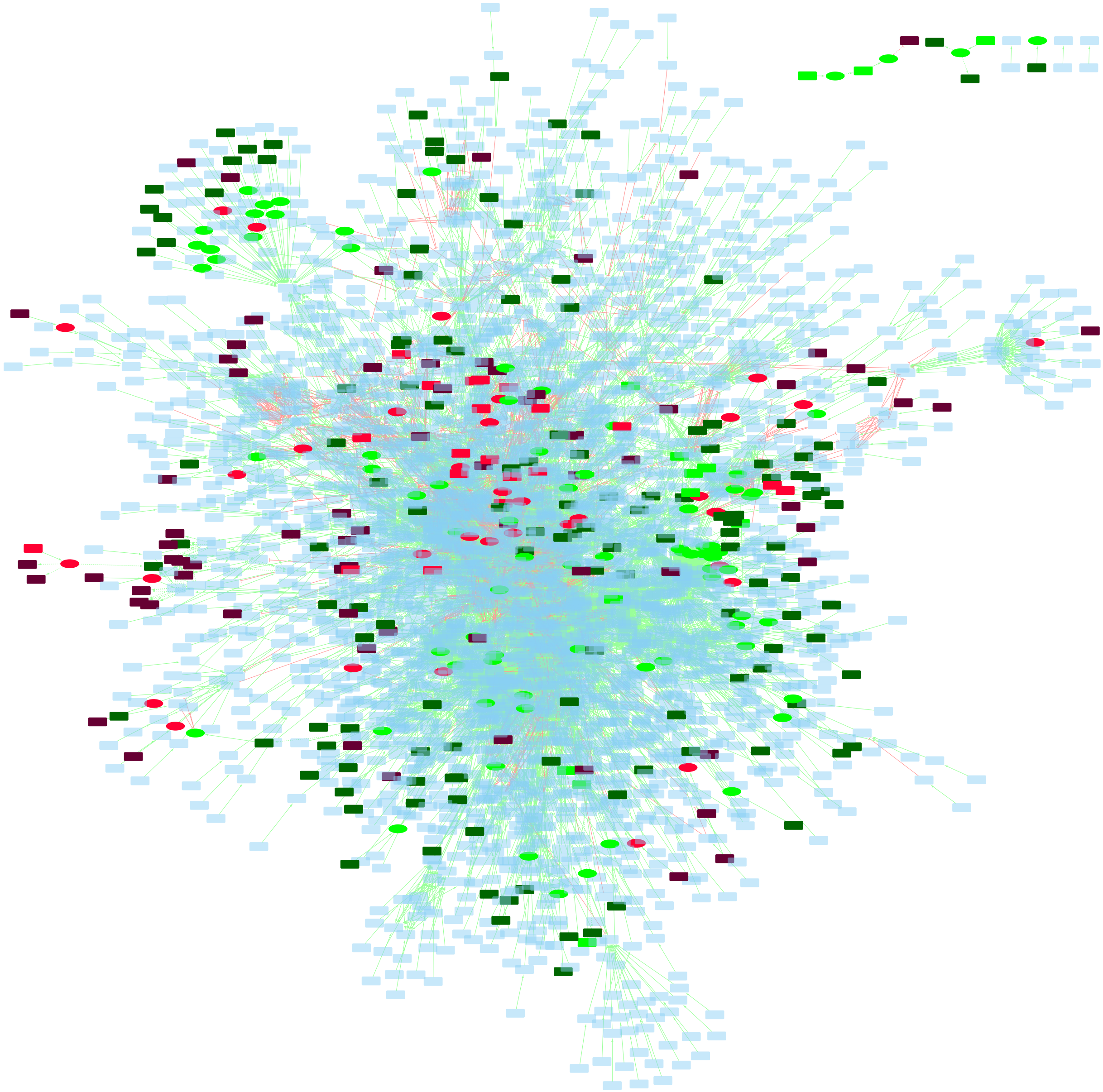

Figure S2: Computational predictions of Iggy in the KEGG graph extraction. The dark green and red nodes are respectively observed as over- and under-expressed; the light green and red nodes are respectively predicted as over- and under-expressed. In the Cytoscape session of Additional file 2: `graph.cys`, this visualisation is available as style `2-Iggy-predictions`.

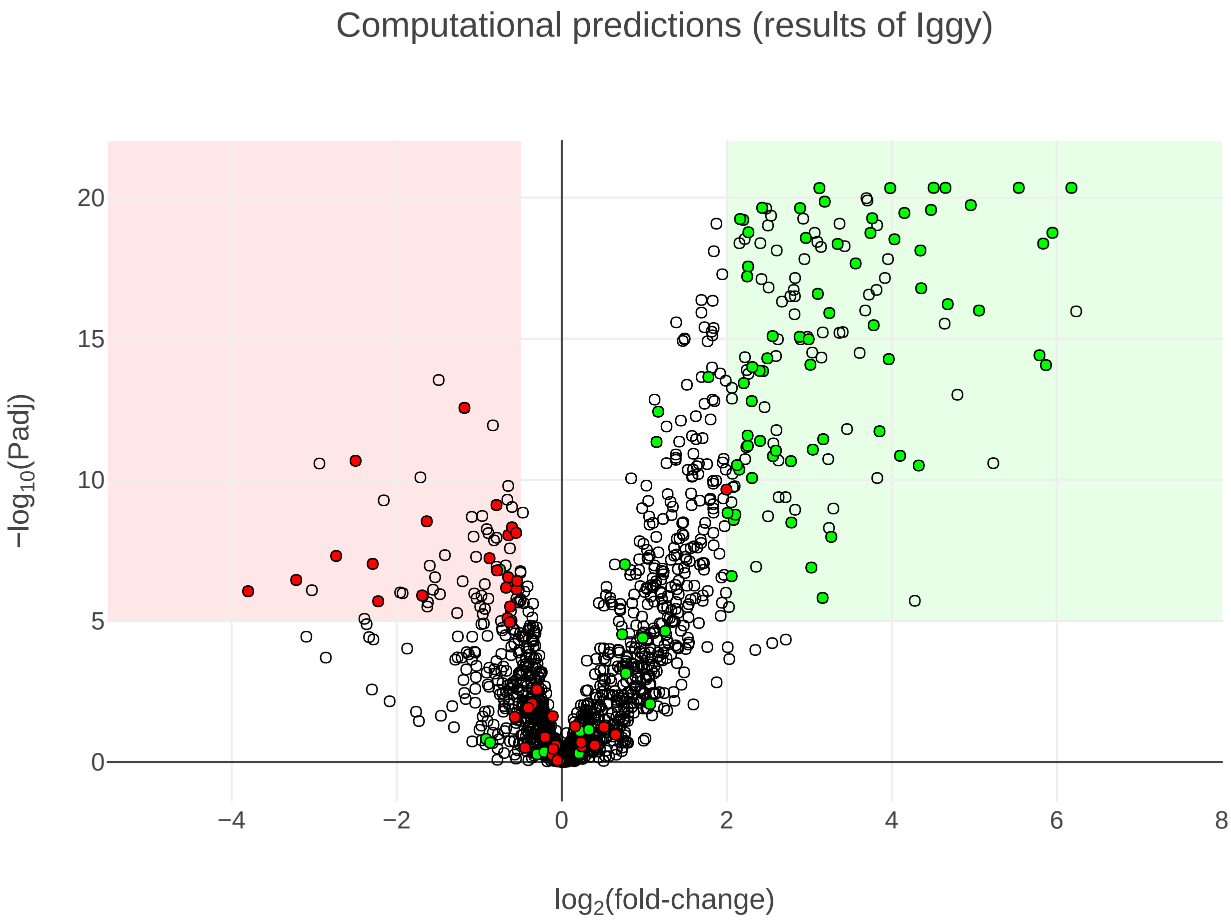

Figure S3: Volcano plot of the genes found in KEGG, given the experimental data from ICGC, with colorings corresponding to the predictions from Iggy. Each dot represents a gene or its corresponding protein in the KEGG graph extraction, plotted regarding its fold-change and P-value from the ICGC experimental data. A green coloured dot is a gene or/and protein predicted up-regulated and a red coloured dot is a gene or/and protein predicted down-regulated. The light red and green areas in the background represent the thresholds beyond which the genes are part of the observations ($\log_2$(fold-change) $< 0.5$ or $\log_2$(fold-change) $> 2$ and $p < 10^{-5}$). An interactive version featuring the names of the genes is available in Additional file 2: `volcano2-predictions.html`.

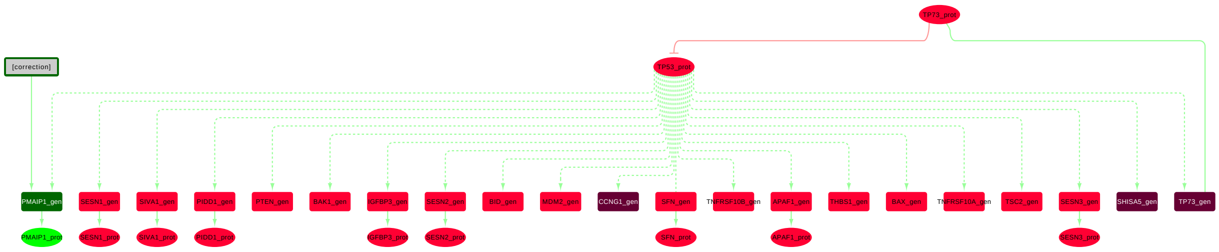

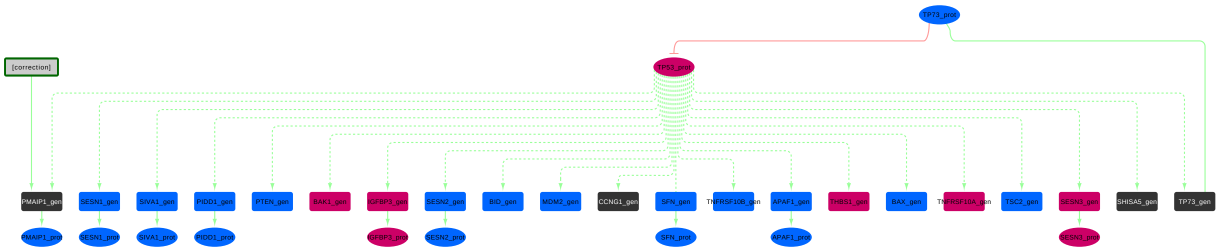

Figure S4: Part of the KEGG extraction graph showing the nodes that are considered unstable regarding their predictions, along with the 4 observed nodes that are downstream of `TP53_prot`. All these unstable nodes happen to be in the neighborhood of `TP53_prot`. Many edges of other incoming and outgoing influences have not been represented in this figure. Furthermore, the minimal correction set (MCoS) repair made in the graph to fix the only inconsistency is reported as a node labeled "`correction`" with an edge towards node `PMAIP1_gen`. Top: the colours match the computational predictions: light green and red nodes are predicted up and down, dark green and red are observed up and down. Bottom: the colours depict the match with the experimental data: the predictions of blue nodes match the experimental data, while the prediction of purple ones do not; black nodes are observations. This graph extract can be found as a network of the Cytoscape session of Additional file 2: `graph.cys`.

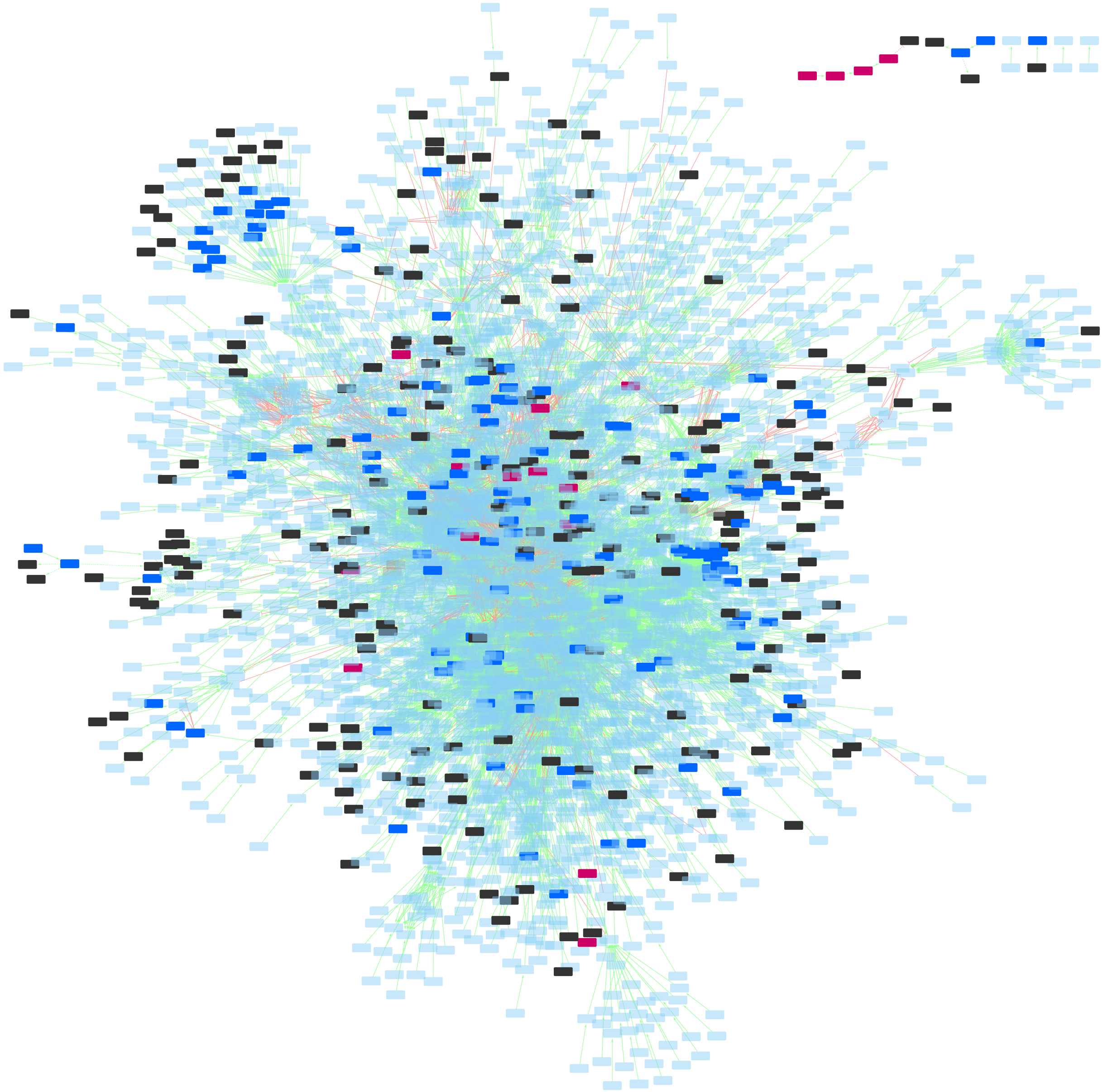

Figure S5: Comparison of the computational predictions of Iggy with the ICGC experimental data. The blue and purple nodes have respectively a matching and non-matching prediction with experimental data; black nodes are the initial observations. In the Cytoscape session of Additional file 2: `graph.cys`, this visualisation is available as style `3-ICGC-comparison`.

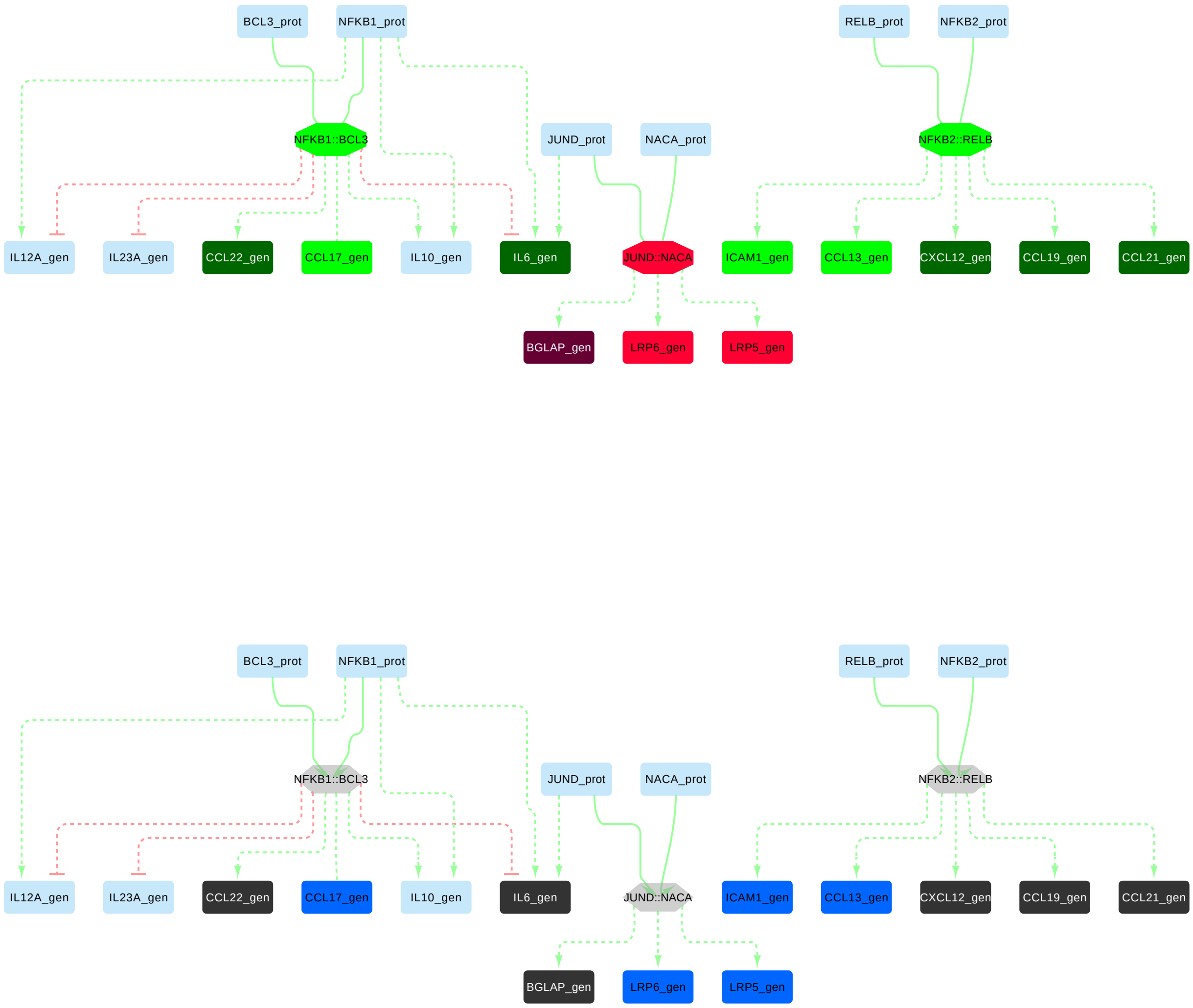

Figure S6: Neighborhood of the predicted complexes `NFKB1::BCL3`, `NFKB2::RELB` and `JUND::NACA`, that bring new information compared to the experimental data. Only the immediate neighbours of these nodes were included; the other upstream and downstream influences of the other nodes are not represented in this figure. Top: the colours match the computational predictions: light green and red nodes are predicted up and down, dark green and red are observed up and down. Bottom: the colours depict the match with the experimental data: the predictions of blue nodes match the experimental data, while the prediction of purple ones do not; black nodes are observations. This graph extract can be found as a network of the Cytoscape session in Additional file 2: `graph.cys`.

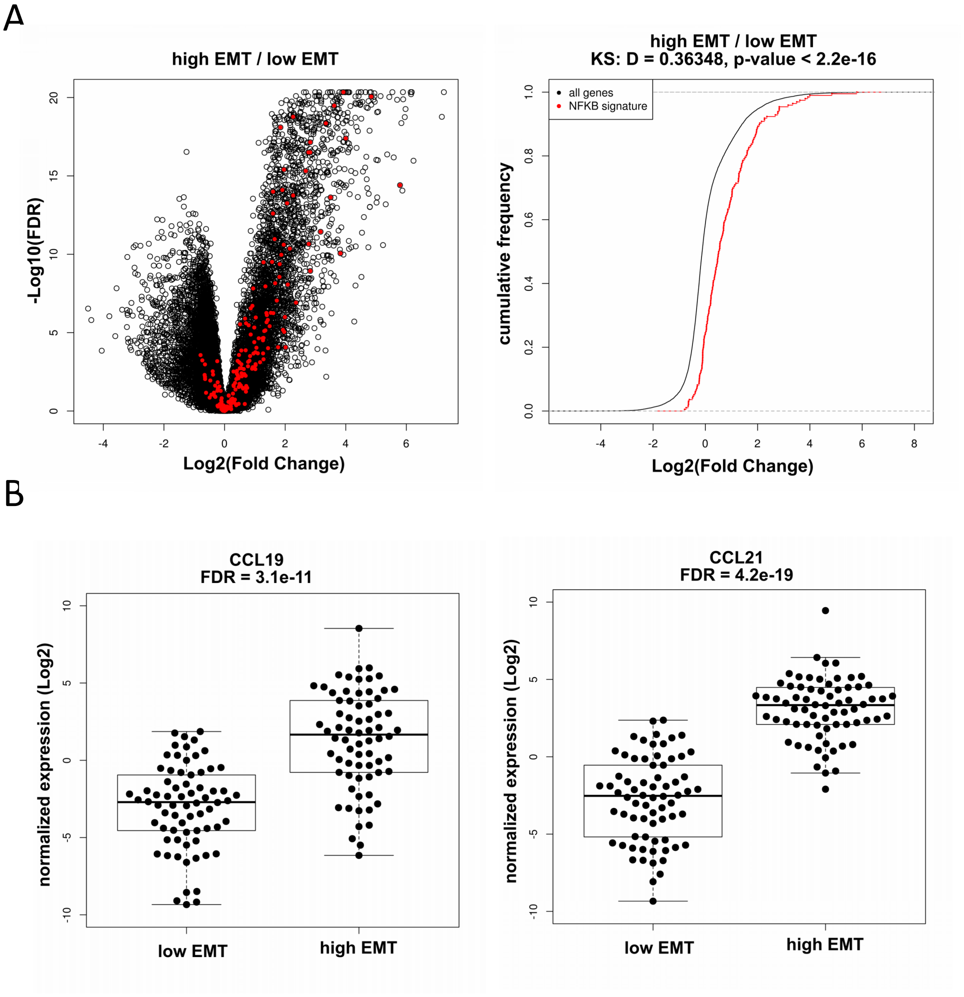

Figure S7: NFKB signaling is activated in aggressive HCC. A) Distribution of gene expression from HALLMARK_TNFA_SIGNALING_VIA_NFKB signature between high and low-EMT HCC. Left panel: volcano plot. Right panel: comparative distribution of NFKB-signature and all genes expressed in HCC. B) Expression of target genes (CCL19 and CCL21) from non canonical NFKB pathways activated by NFKB2-RELB complexes.

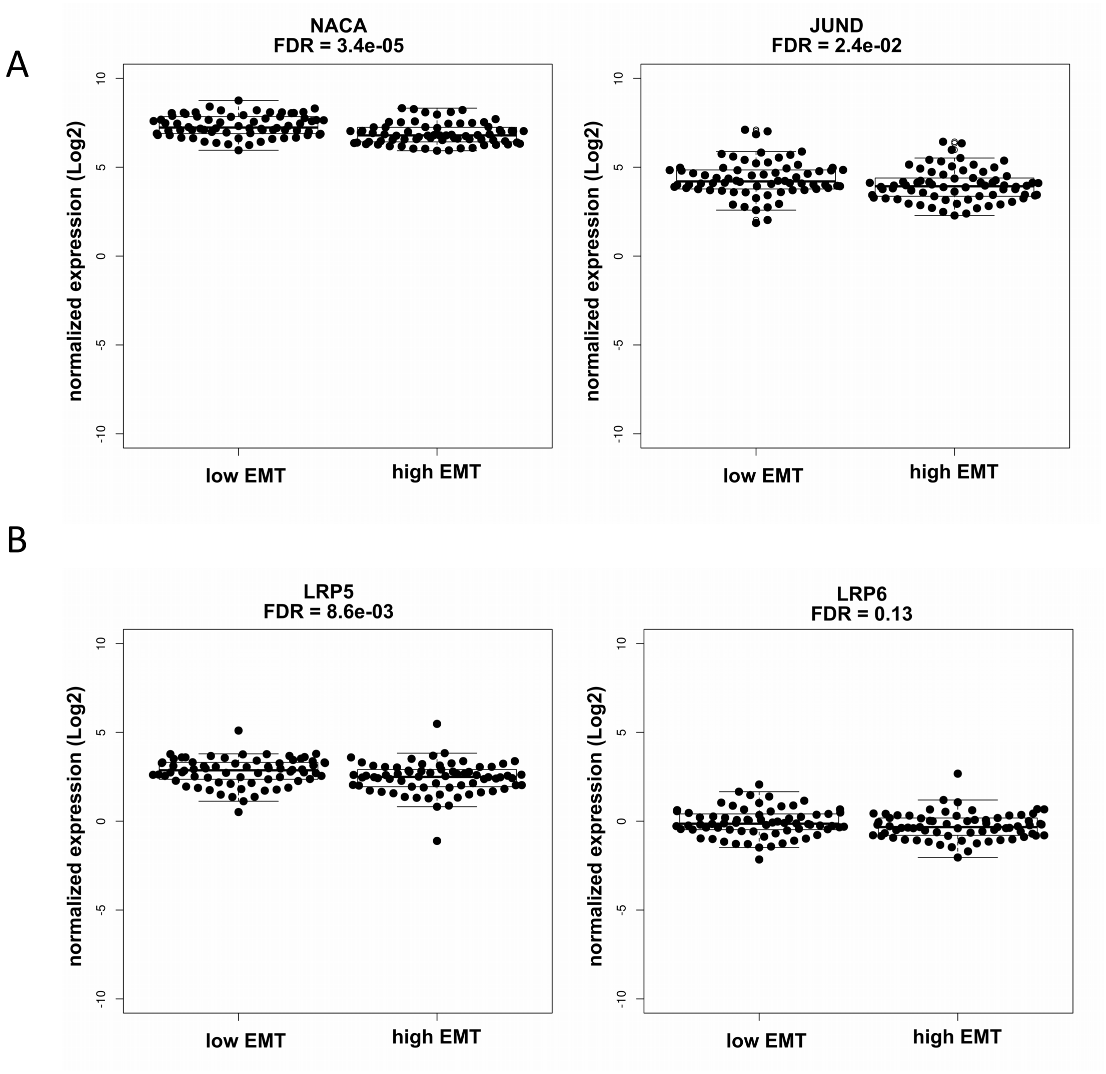

Figure S8: JUND-NACA complex is downregulated in aggressive HCC. A) Distribution of NACA and JUND gene expression between aggressive and non-aggressive HCC. B) Expression of target genes (LRP5 and LRP6) regulated by JUND-NACA complex.

| List of positive observations (up-regulated genes) |
| --- |
| MMP7_gen, DCN_gen, COMP_gen, SFRP5_gen, CCL21_gen, CXCL6_gen, THBS2_gen, KRT19_gen, CXCL14_gen, LAMA2_gen, SLC34A2_gen, CCL11_gen, COL1A1_gen, PDGFRA_gen, COL3A1_gen, SEMA3C_gen, LAMC2_gen, SFRP4_gen, CCL19_gen, FXYD2_gen, EPHA3_gen, SCTR_gen, SLIT2_gen, COL1A2_gen, HHIP_gen, WNT2_gen, NTRK2_gen, CCL26_gen, CXCL1_gen, MMP2_gen, NGFR_gen, ADRA2A_gen, LAMA1_gen, SFRP1_gen, LPAR1_gen, GLI2_gen, ITGA11_gen, CREB3L1_gen, ITGB8_gen, DKK2_gen, EFNA5_gen, ID4_gen, ADCY5_gen, SCD5_gen, PLN_gen, TNC_gen, TRPV6_gen, SFRP2_gen, LAMC3_gen, WNT4_gen, ITGB6_gen, PTGIR_gen, LIF_gen, EPHB3_gen, PPP2R2C_gen, TIMP1_gen, PTPN13_gen, COL6A3_gen, PTH1R_gen, GABBR1_gen, ITGA3_gen, PTHLH_gen, MAPK10_gen, CXCL5_gen, CXCL12_gen, HGF_gen, BHLHE41_gen, EFNB3_gen, ITGA9_gen, PLAT_gen, DHH_gen, COL6A2_gen, FHL2_gen, IL7R_gen, CCL2_gen, EGR2_gen, APLNR_gen, TEK_gen, RASAL1_gen, IL6_gen, PTGS2_gen, ARHGEF4_gen, IGF1R_gen, BMP5_gen, CRYAB_gen, BMPR1B_gen, FGFR1_gen, TGFB2_gen, FZD10_gen, TGFA_gen, NPY1R_gen, NTF3_gen, PRKG1_gen, TGFB3_gen, FZD2_gen, PLXNB3_gen, EDNRA_gen, BDKRB2_gen, F2R_gen, PFKP_gen, CCL22_gen, GLI3_gen, MYL9_gen, NOTCH3_gen, NRG3_gen, FGF1_gen, OLR1_gen, WTIP_gen, FPR1_gen, NTRK3_gen, JAG1_gen, PFKFB3_gen, COL6A1_gen, PTPRR_gen, IL34_gen, CTSK_gen, WNT2B_gen, PLXNA4_gen, F2RL3_gen, PLCB4_gen, THBD_gen, TNXB_gen, COL4A2_gen, CTBP2_gen, TMEM173_gen, DUSP4_gen, HTR2B_gen, FGF18_gen, GDF6_gen, COL4A3_gen, FZD7_gen, OXTR_gen, TGFB1_gen, EGR3_gen, PTGER1_gen, WNT10A_gen, FCER1A_gen, PMAIP1_gen |

| List of negative observations (down-regulated genes) |
| --- |
| CDC23_gen, MAVS_gen, SEC13_gen, CRTC2_gen, SHISA5_gen, RXRB_gen, EIF2B5_gen, RPS6KB2_gen, SENP2_gen, RAF1_gen, PPP2R5D_gen, CCNG1_gen, ACVR2B_gen, RAD9A_gen, FAF1_gen, EIF2B4_gen, ANAPC2_gen, CSNK2B_gen, PPP2R5A_gen, RPTOR_gen, THEM4_gen, CDC26_gen, EIF4EBP2_gen, PHLPP1_gen, DIAPH1_gen, ACACA_gen, SLC38A9_gen, DBI_gen, NPRL2_gen, ELMO1_gen, NR1H3_gen, RXRA_gen, CREB3L4_gen, PPARA_gen, GALT_gen, ACAA1_gen, ANAPC11_gen, SOD1_gen, ERBB3_gen, SMO_gen, THRB_gen, CAT_gen, IRS1_gen, BNIP3_gen, RFNG_gen, BGLAP_gen, FASN_gen, FBXO43_gen, CDC25C_gen, FOXA2_gen, ACSL5_gen, RORC_gen, PLIN5_gen, CD36_gen, CALML6_gen, THPO_gen, ADRB2_gen, TP73_gen, RAC3_gen, ACOX2_gen, SLCO1A2_gen, PROC_gen, THBS4_gen, CCL15_gen, REN_gen, CHAD_gen, SPDYC_gen, TF_gen, APOA2_gen, CCL16_gen, DKK4_gen |

Table S1: List of all observations. All these genes were given as inputs to Iggy.

| Name | Prediction | Fold-change |
|---|---|---|
| NR0B2_gen | + | −0.92 |
| NR0B2_prot | + | −0.92 |
| NR1H4_gen | + | −0.87 |
| NR1H4_prot | + | −0.87 |
| EIF4EBP2_prot | + | −0.75 |
| BMP4_gen | + | −0.30 |
| NR3C2_gen | + | −0.21 |
| NR3C2_prot | + | −0.21 |
| CREB1_prot | - | 0.16 |
| TNFRSF10A_gen | - | 0.23 |
| BAK1_gen | - | 0.24 |
| IGFBP3_gen | - | 0.40 |
| IGFBP3_prot | - | 0.40 |
| TP53_prot | - | 0.51 |
| SESN3_gen | - | 0.65 |
| SESN3_prot | - | 0.65 |
| THBS1_gen | - | 2.00 |

Table S2: List of predictions that are incoherent with the expression data of ICGC. The suffix `_gen` or `_prot` respectively mean that the node models a gene or a protein. The colors emphasize the couples of a gene and the protein corresponding to this gene.

| **List of stable predictions** |
|---|
| SFRP1_prot, LAMA2_prot, VDR_prot, COL4A3_prot, NRG3_prot, VDR_gen, CXCL14_prot, RASAL1_prot, FOXO3_prot, EIF2B5_prot, THEM4_prot, NFKB2::RELB, SCTR_prot, HGF_prot, CCL13_gen, CXCL12_prot, CCL13_prot, THBS2_prot, SFRP4_prot, FGF18_prot, CCL21_prot, ICAM1_gen, CCL19_prot, COL6A3_prot, PHLPP1_prot, PRKG1_prot, HTR2B_prot, PTGIR_prot, FHL2_prot, FGF1_prot, NTF3_prot, TGFA_prot, COL3A1_prot, CXCL6_prot, TNXB_prot, BDKRB2_prot, SENP2_prot, CREB1_prot, FPR1_prot, IL6_prot, PTHLH_prot, CHAD_prot, PPP2R5D_prot, THPO_prot, JAG1_prot, LAMC2_prot, SREBF1_gen, SREBF1_prot, PTGER1_prot, HHIP_prot, GLI3_prot, HIF1A_prot, LIF_prot, LRP6_gen, JUND::NACA, EIF2B4_prot, RAD9A_prot, LRP5_gen, SPDYC_prot, PTPRR_prot, LAMA1_prot, PPP2R2C_prot, COL1A1_prot, RXRB_prot, CCL15_prot, COL6A1_prot, SFRP2_prot, TNC_prot, SGK1_gen, NR3C2_prot, NR3C2_gen, KRAS_gen, COL1A2_prot, PPP2R5A_prot, CTBP2_prot, SEMA3C_prot, CTSK_prot, COL6A2_prot, DKK2_prot, IL34_prot, CCL11_prot, EPHA3_prot, SLC38A9_prot, SLIT2_prot, COL4A2_prot, DCN_prot, LAMC3_prot, THBS4_prot, COMP_prot, DUSP4_prot, WTIP_prot, NOTCH1_gen, NOTCH2_gen, NOTCH4_gen, THRA_prot, BMP4_gen, DKK4_prot, CCL26_prot, NR0B2_prot, NR1H4_gen, NR0B2_gen, NR1H4_prot, SFRP5_prot, CCL16_prot, NFKB1::BCL3, CCL17_prot, CCL17_gen, CCL22_prot, CSNK2B_prot, RFNG_prot, ADRA2A_prot, ELMO1_prot, EFNB3_prot, NFATC1_prot, CXCL5_prot, NTRK3_prot, EIF4EBP2_prot, PTH1R_prot |

| **List of unstable predictions** |
|---|
| PIDD1_prot, BAK1_gen, THBS1_gen, SESN3_prot, SESN3_gen, APAF1_gen, TP73_prot, TSC2_gen, PTEN_gen, SESN1_prot, SIVA1_prot, SESN2_prot, APAF1_prot, SIVA1_gen, SESN1_gen, TNFRSF10A_gen, SFN_gen, IGFBP3_prot, BID_gen, BAX_gen, SFN_prot, IGFBP3_gen, TNFRSF10B_gen, MDM2_gen, PIDD1_gen, TP53_prot, SESN2_gen, PMAIP1_prot |

Table S3: List of stable and unstable predictions, based on the stability study.

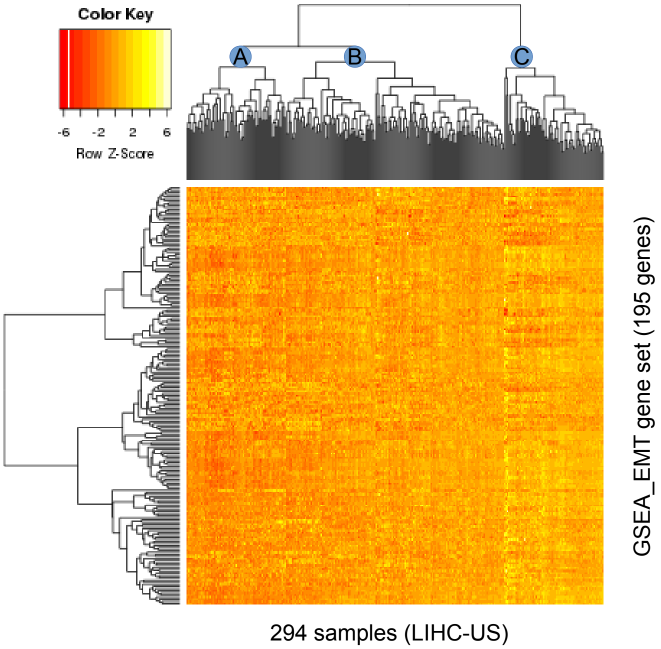

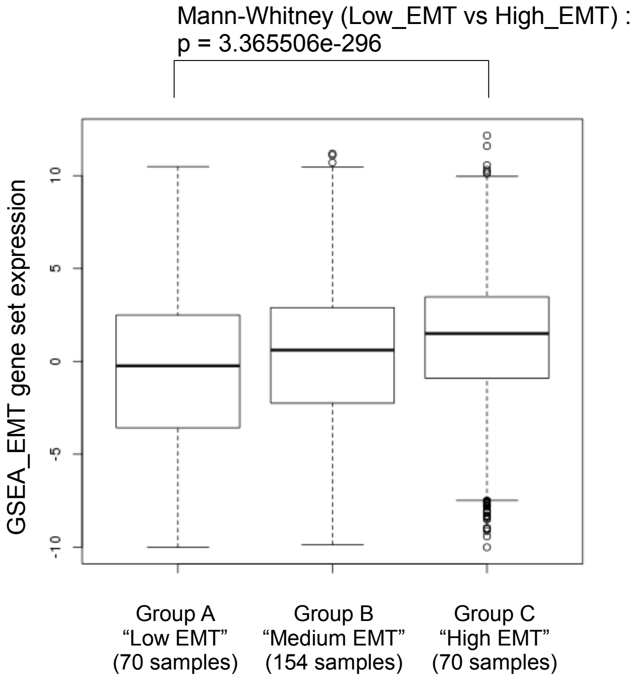

Figure S9: Top: Expression heat map of EMT signature genes (195 genes after removal of undetectable genes) on the 294 experimental samples (from LIHC-US project of ICGC, with one sample per subject). Above the heat map is featured the hierarchy provided by the clustering analysis, where the letters A, B and C represent the three main groups that are identified with this method. Bottom: Expression of EMT signature genes averaged on all subjects for each group returned by the clustering method.

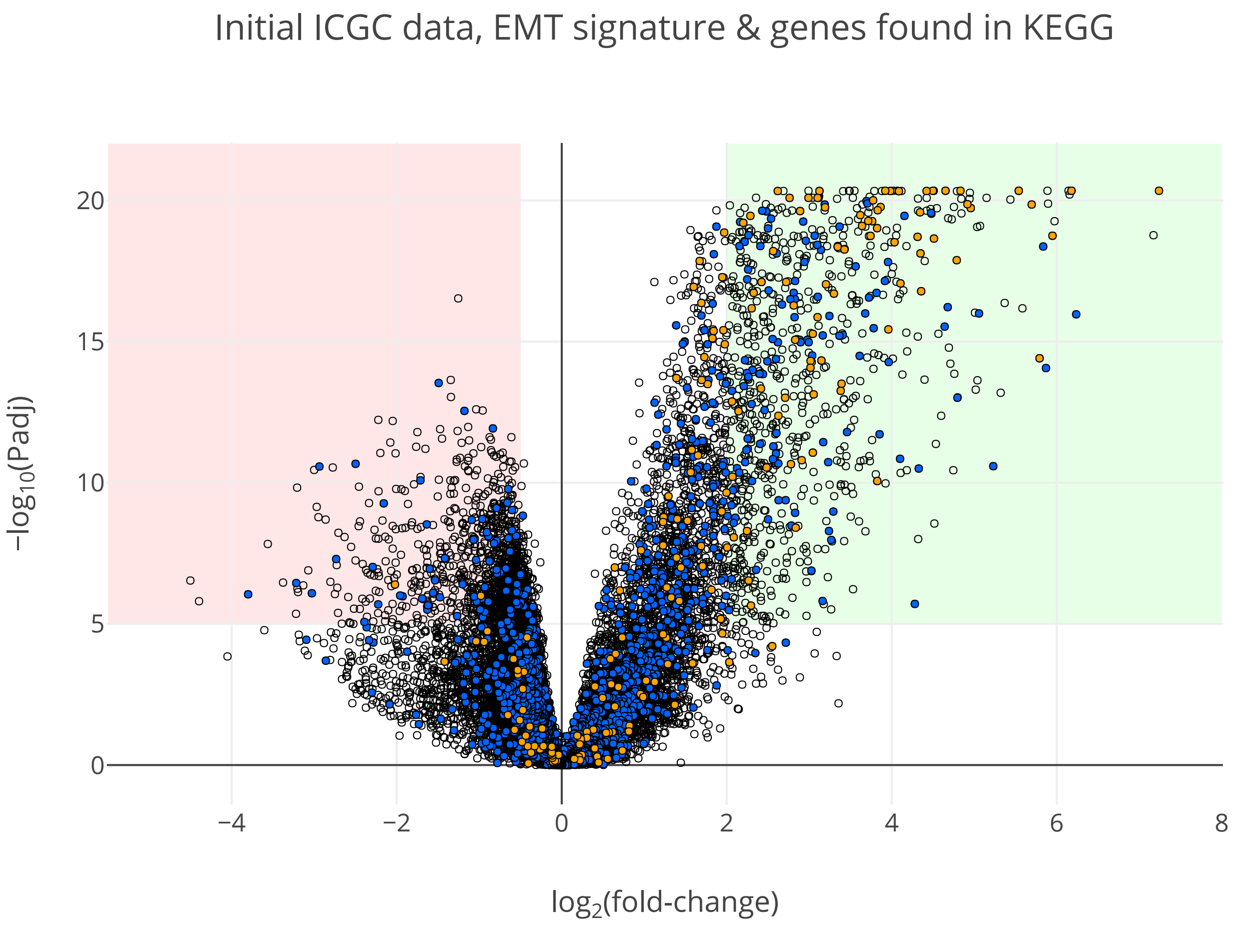

Figure S10: Volcano plot of the experimental data extracted from ICGC, highlighting the EMT signature and the result of the KEGG extraction. Each circle represents a gene from the ICGC data. The genes of the EMT signature are filled in orange while genes that were found in the KEGG extraction are filled in blue. The green and red background areas highlight the sets of genes that are considered as respectively over- and underexpressed in the present work. An interactive version featuring the names of the genes is available in Additional file 2: `volcano1-all-genes.html`.

### Building the Signaling Network from the KEGG Pathway Database

This appendix gives an in-depth description of how the human signaling network was built from the KEGG Pathway database. It is the full explanation of what was summarized in Section 5.2 of the main article.

#### 1 Using the KEGG Pathway database

This work was performed on a human signaling network derived from the KEGG Pathway database. This database mostly contains metabolic and signaling networks for a couple of species, including *homo sapiens*. In this work, only human signaling networks were considered. KEGG Pathway is divided into seven sections:

1. Metabolism
2. Genetic information processing
3. Environmental information processing
4. Cellular processes
5. Organismal systems
6. Human diseases
7. Drug development

The section 1 contains the metabolic networks. The section 7 is somewhat apart: it contains drug classifications as well as synthesis pathways. All the other sections contain the signaling networks.

All the human signaling pathways of the sections 2, 3, 4 and 5 were fetched from KEGG Pathway using its API. Each of these pathways is encoded in its native file format: the KGML (KEGG Markup Language). Note that the section 6 was excluded even if it also contains human signaling pathways. As its name indicates, this section implements specific pathological features. However, the goal was to obtain a generic human signaling network independently of specific features such as diseases and mutations.

Once fetched, the KGML files were converted to the SIF file format (Simple Interaction Format), a TSV file format useful when working with networks because each line encodes an edge in an intuitive way:

source `\t` relation `\t` target

where `\t` stands for a horizontal tab character. The signaling pathways converted to the SIF file format were then merged into one file to obtain a generic human signaling network. Because the data used in the present work are about gene expression levels, a clear distinction was made between nodes representing genes and nodes representing gene products, namely proteins.

#### 2 Distinguishing genes and their products

In the KEGG Pathway database, the distinction between genes and their products is implicit: nodes can either represent proteins or their corresponding genes. This information is embedded in the relation types, particularly `PPrel` edges (protein-protein relations) and `GErel` edges (gene expression relations). `PPrel` edges indicate that both the source and target nodes are proteins. `GErel` edges indicate that source nodes are transcription factors and that target nodes are genes. Therefore, to explicitly differentiate genes and proteins, the source nodes of `GErel` edges were suffixed with `.prot` to indicate proteins, and the target nodes were suffixed with `.gen` to indicate genes. Concerning `PPrel` edges, both the source and target nodes were suffixed with `.prot`.

Furthermore, in order to link genes and their products, a relation type was added: the `GPrel` type (gene-protein relations). `GPrel` edges indicate that genes can produce their corresponding products. Consequently, for each node modeling a gene, a `GPrel` edge starting from it and ending on an added node suffixed with `.prot` was added. These added nodes therefore model the corresponding gene products while the `GPrel` edges model the gene expressions themselves.

Altogether, the human signaling network can now explicitly model protein-protein interactions, gene expression regulations, and gene expressions as illustrated in Figure S11 of this Appendix and Figure S12 of this Appendix. Note that a clear distinction between the regulation of gene expression (`GErel` edges) and gene expression itself (`GPrel` edges) was consequently implemented in the network.

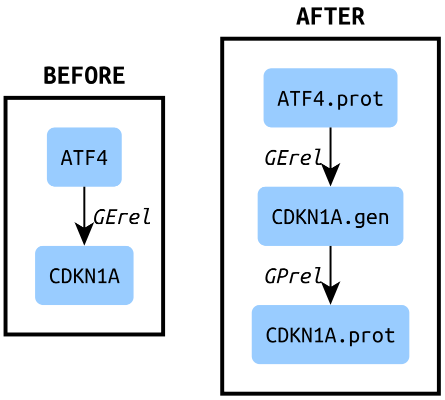

Figure S11: As an example, this figure shows the interaction from the transcription factor ATF4 to the target gene CDKN1A. The *before* box represents how it is in KEGG Pathway: the edge is of `GErel` type, implicitly meaning that the source node ATF4 is a transcription factor and that the target node CDKN1A is a gene. The *after* box shows how it is after explicitness: node types are indicated by suffixes, a node is added to model the gene product, and a `GPrel` edge is added to link the gene and its product, thus modeling gene expression apart from its regulation. Consequently, the gene and its product become two distinct entities involved in distinct relations.

Finally, in order to allow the genes concerned by the data used in the present work to match their corresponding nodes in the human signaling network, a `GPrel` edge was added for each node. Except when already done due to a `GErel` edge as explained above, each node implicitly modeling a protein was put as target of a `GPrel` edge with the suffix `.prot`. By doing so, a source node was added for each of these `GPrel` edges with the suffix `.gen`: these are the corresponding genes, possibly concerned by the used data, as illustrated in Figure S12 of this Appendix.

#### 3 Selecting functional interactions

In the KEGG Pathway database, in addition to their relation types such as `PPrel` or `GErel`, each edge can be annotated with one ore more keywords bringing details about the modeled interaction. Four of these keywords explicitly specify the sign of the interactions, that is, if an influence is positive/activating or negative/inhibiting. These four functional keywords are “activation” and “inhibition” for `PPrel` edges, and “expression” and “repression” for `GErel` edges. The other keywords can not be used to systematically infer edge signs. For example, the keyword “phosphorylation” can be present in positive and negative edges because the functional impact of phosphorylating the target depends on the target itself.

The human signaling network used in the present work needs to be signed. As explained in Section 5.3 of the main article, a next step consists in running the predicting tool Iggy in order to infer the state of unobserved nodes using observed nodes and logical rules. The observed nodes are genes for which the ICGC experimental data gives an information about over- or under-expression between aggressive and non-aggressive HCC. The unobserved node are the remaining nodes of the network, namely the genes devoid of such data together with all the proteins (because experimental data are about gene expressions, not about protein activities). Because the logical rules implemented in Iggy allow to infer the state of a given node depending on the state of its predecessors and successors, and according to the sign of the edges linking them, edges have to be signed (positive and negative influences only) in our model.

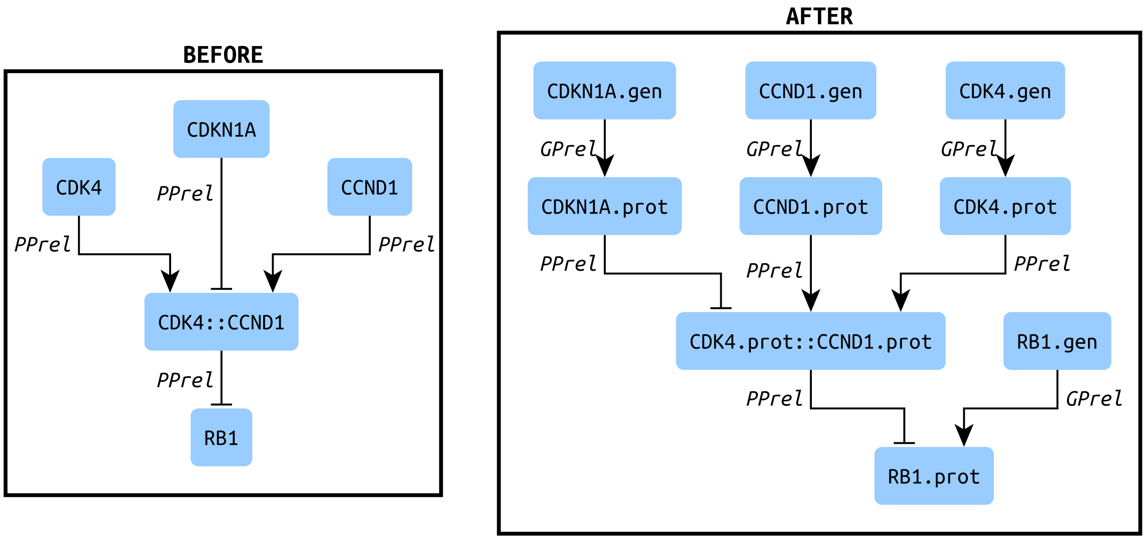

Figure S12: As an example, this figure depicts the formation of the CDK4-CCND1 complex, its possible inhibition by CDKN1A and its inhibiting effect on RB1. The *before* box shows how it is in KEGG Pathway: the edges are of `PPrel` type, implicitly meaning that the interacting nodes are proteins. The *after* box shows how it is after explicitness: node types are indicated by suffixes, each node modeling a protein is put as target of a `GPrel` edge, and nodes are added to model the source genes. Therefore, gene expression data can be injected into the network without ambiguity. Complexes are named after the list of their component names separated by "`::`".

Note that using this approach on a network where nodes modeling genes and nodes modeling proteins are distinct allows to predict protein states from data about gene expressions. It can be insightful because the final effectors of biological functions are proteins, not genes: genes encode information but proteins perform the work. However, obtaining large scale data about protein activities is more challenging than obtaining large scale data about gene expressions. Consequently, such a predicting approach is particularly interesting, especially because the expression of a gene does not systematically imply that the produced protein is functional.

To obtain the signed and functional human signaling network, only the edges bringing one of the four functional keywords mentioned above were selected. Once done, edge signs were inferred and annotated according to the syntax required by Iggy: 1 for positive edges and $-1$ for negative ones. Moreover, special characters in nodes names, such as spaces and dots (`.`), were replaces by underscores (_).

#### 4 Extracting regulatory signaling pathways

Once the signed human signaling network obtained, the signaling pathways regulating the genes of interest were extracted. These regulatory signaling pathways are the upPathrider paths of the nodes modeling the genes of interest. The tool *Pathrider* was developed and used in this context, with a command *Stream* especially designed for the purpose of extracting upstream paths. Pathrider is freely available on GitHub at `https://github.com/arnaudporet/pathrider` under the Simplified BSD License. It is on these extracted signaling pathways that predictions about node states using Iggy were performed.

Given a network, Pathrider performs upstream or downstream pathfinding starting from a list of root nodes up to terminal nodes, namely nodes with no incoming edges (indegree = 0) in the upstream case and nodes with no outgoing edges (outdegree = 0) in the downstream case. Here the network was the human signaling network derived from KEGG Pathway, and upstream pathfinding was performed with our list of 1913 differentially expressed genes as root nodes (see Section 5.1 of the main article).

Forthermore, Pathrider has a blacklist option allowing to prevent the exploration of a given list of biomolecules during the upPathrider exploration. In our case, this option has been used with the list of 4220 gene names whose expression is undetectable (see Section 5.1 of the main article) to avoid including genes and proteins having one of these names.

A final supplementary step consisted in also filtering out the protein complexes containing one of these names afterwards.

| | | Complete network (with `GPrel`) | Network without `GPrel` |
|---|---|---|---|
| | **Number of observations** | 209 | 63 |
| | **Number of predictions** | 148 | 23 |
| strong | **positive** (+) | 92 | 13 |
| strong | **negative** (−) | 54 | 5 |
| strong | no-change (0) | 0 | 3 |
| weak | may-up (NOT−) | 0 | 0 |
| weak | may(down (NOT+) | 0 | 0 |
| weak | change (CHANGE/NOT0) | 2 | 2 |
| | Found in ICGC data | 141 | 18 |
| | Coherent with ICGC data | 124 (88%) | 9 (50%) |

Table S4: Summary of Iggy's predictions when `GPrel` edges are included in (left column) or excluded from (right column) the network. The number of observed (input) nodes also varies because the removal of `GPrel` edges also removes nodes that are not connected to other nodes anymore.

| **List of positive (up-regulated) predictions** |
|---|
| `CHUK_prot`, `EIF4EBP2_prot`, `GLI3_prot`, `HIF1A_prot`, `NFATC1_prot`, `NFKB1::BCL3`, `NFKB2_prot`, `NFKB2::RELB`, `NR0B2_prot`, `NR3C2_prot`, `RELB_prot`, `THRA_prot`, `VDR_prot` |

| **List of negative (down-regulated) predictions** |
|---|
| `CREB1_prot`, `FOXO3_prot`, `JUND::NACA`, `SREBF1_prot`, `TP53_prot` |

Table S5: List of positive and negative predictions returned by Iggy on the network without `GPrel` edges. The predictions that are common with the predictions on the complete network (containing `GPrel` edges) are coloured in blue.
