## Additional information to generate differentially expressed genes from ICGC database for "A pipeline to create predictive functional networks: application to the tumor progression of hepatocellular carcinoma": dataset_filtering.sh

```
#!/bin/bash

# Dataset filtering
# -----------------
# This file is part of the Supplementary Material of the submission entitled:
# A pipeline to create predictive models: application to the tumor progression of hepatocellular carcinoma
# Authors: Maxime Folschette, Vincent Legagneux, Arnaud Poret, Lokmane Chebouba, Carito Guziolowski and Nathalie Théret

# This script filters data from an ICGC dump, to keep only expression data of samples in project
# LIHC-US and corresponding to primary tumor.
# Usage: Called from script donwnload-and-run.sh, or:
#   bash dataset_filtering.sh

# Inputs:
# - exp_seq.LIHC-US.tsv downloaded from https://dcc.icgc.org/api/v1/download?fn=/release_21/Projects/LIHC-US/exp_seq.LIHC-US.tsv.gz
# - specimen.LIHC-US.tsv downloaded from https://dcc.icgc.org/api/v1/download?fn=/release_21/Projects/LIHC-US/specimen.LIHC-US.tsv.gz
# - sample.LIHC-US.tsv downloaded from https://dcc.icgc.org/api/v1/download?fn=/release_21/Projects/LIHC-US/sample.LIHC-US.tsv.gz

### Retrieving ICGC data
## https://dcc.icgc.org/
# download date: 2016/07/19
# select datasets with RNAseq expression data (EXP-S data type)
# retrieve Sequencing-based Gene Expression data:
# exp_seq.tsv.gz file

# Files:
# exp_seq.tsv

## https://dcc.icgc.org/
# retrieve clinical data:
# icgc-dataset-1468914478976.tar

# Files:
# donor_family.tsv
# specimen.tsv
# donor_therapy.tsv
# sample.tsv
# donor_exposure.tsv
# donor.tsv

### File names
FILE_EXPSEQ="exp_seq.LIHC-US.tsv"
FILE_SPECIMEN="specimen.LIHC-US.tsv"
FILE_SAMPLE="sample.LIHC-US.tsv"

### Selecting expression data from LIHC-US project (TCGA Liver Hepatocarcinoma)
## Bash
head -1 "$FILE_EXPSEQ" > exp_seq_headers.tsv
grep -w 'LIHC-US' "$FILE_EXPSEQ" > exp_seq_LIHC-US.tsv

### Simplifying data
## Bash
cut -f2,4,8,9 exp_seq_headers.tsv > exp_seq_headers_simple.tsv
cut -f2,4,8,9 exp_seq_LIHC-US.tsv > exp_seq_LIHC-US_simple.tsv
# visualizing selected fields:
sed 's/\t/\n/g' exp_seq_headers_simple.tsv
# project_code
# icgc_sample_id
# gene_id
# normalized_read_count

### Selecting LIHC specimens corresponding to primary solid tumor
## Bash
grep -w 'LIHC-US' "$FILE_SPECIMEN" | grep 'Primary\ tumour\ \-\ solid\ tissue' | cut -f1 > Primary_Tumor_Specimens.txt
### Selecting samples corresponding to these specimens
## Bash
for i in $(cat Primary_Tumor_Specimens.txt); do grep -w $i "$FILE_SAMPLE"; done | cut -f1 >> Primary_Tumor_Samples.txt
### Selecting expression data corresponding to these samples
## Bash
#for i in $(cat Primary_Tumor_Samples.txt); do grep -w $i exp_seq_LIHC-US_simple.tsv >> exp_seq_LIHC-US_simple_Primary_Tumor.tsv; done
grep -w -f Primary_Tumor_Samples.txt exp_seq_LIHC-US_simple.tsv >> exp_seq_LIHC-US_simple_Primary_Tumor.tsv

### Checking
## Bash
cut -f2 exp_seq_LIHC-US_simple.tsv | sort -u | wc
#     345     345    3105
# 345 RNAseq samples in LIHC-US
cut -f2 exp_seq_LIHC-US_simple_Primary_Tumor.tsv | sort -u | wc
#     294     294    2646
# 294 RNAseq samples in LIHC-US restricted to primary solid tumors
```
