## Additional information to generate differentially expressed genes from ICGC database for "A pipeline to create predictive functional networks: application to the tumor progression of hepatocellular carcinoma": diffexp_and_clustering.R

```
# Clustering and differential expression analysis
# -----------------------------------------------
# This file is part of the Supplementary Material of the submission entitled:
# Hepatocellular carcinoma computational models identify key protein-complexes associated to tumor progression
# Authors: Maxime Folschette, Vincent Legagneux, Arnaud Poret, Carito Guziolowski and Nathalie Théret

# This script applies the differential analysis to the experimental gene expression data,
# then the clustering analysis, and finally ouputs files and plots.
# Usage: Called from script donwnload-and-run.sh, or:
#   bash dataset_filtering.sh

library("gplots")
library(RColorBrewer)
mypalette=colorRampPalette(c("blue","white","red"))(n = 21)

## Retrieving LIHC whole data
# Depending on file location:
#whole=read.table("exp_seq_LIHC-US_simple_Primary_Tumor.tsv",sep="\t",header=FALSE)
whole=read.table("exp_seq_LIHC-US_simple_Primary_Tumor.tsv",sep="\t",header=FALSE)
colnames(whole)=c("project_code","icgc_sample_id","gene_id","normalized_read_count")

## remove unused levels
whole=droplevels(whole)

## generate a two-entry table (samples_x_genes) of expression values (norm.allgenesalized counts per gene)
whole.tab=tapply(whole$normalized_read_count,list(whole$gene_id,whole$icgc_sample_id), function(x) as.numeric(x))

## extract numeric values
whole.mat=apply(whole.tab,c(1,2),function(x) x[[1]][1])
dim(whole.mat)
# [1] 20502   294

## log values (add 1e-8, a very low value based on distribution, to avoid infinite log values)
whole.log=log(whole.mat+1e-8, 2)

## detect not/weakly expressed genes
sample.med=apply(whole.log,1,median)
hist(sample.med, breaks=50)

# => cut-off at -25
## delete entries corresponding to not/weakly expressed genes
whole.log=whole.log[sample.med>(-25),]
dim(whole.log)
# [1] 16282   294

### Normalization
# centering data on median
whole.med=apply(whole.log,2,median)
whole.norm=sweep(whole.log,2,whole.med,"-")

## Retrieving "GSEA_EMT" signature

# http://software.broadinstitute.org/gsea/msigdb/cards/HALLMARK_EPITHELIAL_MESENCHYMAL_TRANSITION.html
# Gene Set: HALLMARK_EPITHELIAL_MESENCHYMAL_TRANSITION
# 200 genes

GSEA_EMT_genes=c("ABI3BP", "ACTA2", "ADAM12", "ANPEP", "APLP1", "AREG", "BASP1", "BDNF", "BGN", "BMP1", "CADM1", "CALD1", "CALU", "CAP2", "CAPG", "CD44", "CD59", "CDH11", "CDH2", "CDH6", "COL11A1", "COL12A1", "COL16A1", "COL1A1", "COL1A2", "COL3A1", "COL4A1", "COL4A2", "COL5A1", "COL5A2", "COL5A3", "COL6A2", "COL6A3", "COL7A1", "COL8A2", "COMP", "COPA", "CRLF1", "CTGF", "CTHRC1", "CXCL1", "CXCL12", "CXCL6", "CYR61", "DAB2", "DCN", "DKK1", "DPYSL3", "DST", "ECM1", "ECM2", "EDIL3", "EFEMP2", "ELN", "EMP3", "ENO2", "FAP", "FAS", "FBLN1", "FBLN2", "FBLN5", "FBN1", "FBN2", "FERMT2", "FGF2", "FLNA", "FMOD", "FN1", "FOXC2", "FSTL1", "FSTL3", "FUCA1", "FZD8", "GADD45A", "GADD45B", "GAS1", "GEM", "GJA1", "GLIPR1", "GLT25D1", "GPC1", "GPX7", "GREM1", "HTRA1", "ID2", "IGFBP2", "IGFBP3", "IGFBP4", "IL15", "IL32", "IL6", "IL8", "INHBA", "ITGA2", "ITGA5", "ITGAV", "ITGB1", "ITGB3", "ITGB5", "JUN", "LAMA1", "LAMA2", "LAMA3", "LAMC1", "LAMC2", "LEPRE1", "LGALS1", "LOX", "LOXL1", "LOXL2", "LRP1", "LRRC15", "LUM", "MAGEE1", "MATN2", "MATN3", "MCM7", "MEST", "MFAP5", "MGP", "MMP1", "MMP14", "MMP2", "MMP3", "MSX1", "MXRA5", "MYL9", "MYLK", "NID2", "NNMT", "NOTCH2", "NT5E", "NTM", "OXTR", "PCOLCE", "PCOLCE2", "PDGFRB", "PDLIM4", "PFN2", "PLAUR", "PLOD1", "PLOD2", "PLOD3", "PMEPA1", "PMP22", "POSTN", "PPIB", "PRRX1", "PRSS2", "PTHLH", "PTX3", "PVR", "QSOX1", "RGS4", "RHOB", "SAT1", "SCG2", "SDC1", "SDC4", "SERPINE1", "SERPINE2", "SERPINH1", "SFRP1", "SFRP4", "SGCB", "SGCD", "SGCG", "SLC6A8", "SLIT2", "SLIT3", "SNAI2", "SNTB1", "SPARC", "SPOCK1", "SPP1", "TAGLN", "TFPI2", "TGFB1", "TGFBI", "TGFBR3", "TGM2", "THBS1", "THBS2", "THY1", "TIMP1", "TIMP3", "TNC", "TNFAIP3", "TNFRSF11B", "TNFRSF12A", "TPM1", "TPM2", "TPM4", "VCAM1", "VCAN", "VEGFA", "VEGFC", "VIM", "WIPF1", "WNT5A")

## Subdata corresponding to the expressions of GSEA_EMT signature genes in LIHC
subdata.emt=whole.norm[rownames(whole.norm) %in% GSEA_EMT_genes,]
dim(subdata.emt)
# [1] 195 294
# 5 genes are undetectable in LIHC samples

### Expression map & clustering on GSEA_EMT gene set
## Expression heat-map of GSEA_EMT genes in LIHC samples
png("ICGC_LIHC_GSEA_EMT_Expression_complete_colorkey.png")
heatmap.2(subdata.emt, distfun=function(x) dist(x, method = "euclidean") ,hclustfun=function(x) hclust(x,method="complete"), scale="row", key=TRUE, symkey=FALSE, density.info="none", trace="none"
)
dev.off()
# This allows identifying 3 groups corresponding to low, medium and high expression of the GSEA_EMT gene set.

## defining 3 sample groups (clusters), based on GSEA_EMT clustering
emt.cut=cutree(hclust(dist(t(subdata.emt), method = "euclidean"),method="complete"), k=3)
grp1=emt.cut[emt.cut==1]
grp2=emt.cut[emt.cut==2]
grp3=emt.cut[emt.cut==3]
# number of samples in each cluster
length(grp1)
# [1] 70
length(grp2)
# [1] 70
length(grp3)
# [1] 154

## defining the "low_EMT", "medium_EMT" and "high_EMT" groups
low.EMT=names(grp1)
high.EMT=names(grp2)
medium.EMT=names(grp3)

## expression of GSEA_EMT genes in these 3 groups
png("boxplot_GSEA_EMT_low_medium_high_EMT.png")
boxplot(as.numeric(subdata.emt[,low.EMT]),as.numeric(subdata.emt[,medium.EMT]),as.numeric(subdata.emt[,high.EMT]), names=c("low_EMT","medium_EMT","high_EMT"))
dev.off()

## comparing expression of GSEA_EMT genes in "low_EMT" and "high_EMT" groups (Mann-Whitney)
wilcox.test(as.numeric(subdata.emt[,low.EMT]),as.numeric(subdata.emt[,high.EMT]))[[3]][1]
# [1] 3.365506e-296

### differential expression analysis of all genes in the "low_EMT" and "high_EMT" groups
## defining sample groups corresponding to "low_EMT" or "high_EMT" groups
LIHC.lowEMT=whole.norm[,colnames(whole.norm)%in%low.EMT]
LIHC.highEMT=whole.norm[,colnames(whole.norm)%in%high.EMT]
## calculating fold-changes (Log2) for each gene in "high_EMT" versus "low_EMT" groups
Log2FC=sapply(rownames(whole.norm), function(x) mean(LIHC.highEMT[x,])-mean(LIHC.lowEMT[x,]))
## calculating p-values (Mann-Whitney) for each gene between "high_EMT" and "low_EMT" groups
pval=sapply(rownames(whole.norm), function(x) wilcox.test(LIHC.highEMT[x,],LIHC.lowEMT[x,])[[3]][1])
## adjusting p-value for multiple analysis (Benjamini & Hochberg)
padj=p.adjust(pval, method="BH")

## volcano plot of all genes with GSEA_EMT genes in red
png("volcano_EMThigh_vs_EMTlow.png")
plot(Log2FC, -log(padj,10))
points(Log2FC[names(Log2FC) %in% GSEA_EMT_genes], -log(padj[names(padj) %in% GSEA_EMT_genes],10), pch=20, col="red")
dev.off()

## output differential expression data
# output whole diffexp data
diffexp=data.frame(names(Log2FC),Log2FC,padj)
diffexp$volcano=diffexp$Log2FC*(-log(diffexp$padj,10))
colnames(diffexp)=c("Genes","Log2FC","padj","volcano")
diffexp=diffexp[order(diffexp$volcano, decreasing=TRUE),]
write.table(diffexp,"GSEA_EMThigh_vs_EMTlow_diffexp.csv", quote=FALSE, sep="\t",row.names=FALSE)

# output diffexp data for the GSEA_EMT gene set
diffexp.EMT=diffexp[diffexp$Genes %in% GSEA_EMT_genes,]
diffexp.EMT=diffexp.EMT[order(diffexp.EMT$volcano, decreasing=TRUE),]
write.table(diffexp.EMT,"GSEA_EMThigh_vs_EMTlow_diffexp_EMT.csv", quote=FALSE, sep="\t",row.names=FALSE)

# output diffexp data for genes not in the GSEA_EMT gene set
diffexp.notEMT=diffexp[!diffexp$Genes %in% GSEA_EMT_genes,]
diffexp.notEMT=diffexp.notEMT[order(diffexp.notEMT$volcano, decreasing=TRUE),]
write.table(diffexp.notEMT,"GSEA_EMThigh_vs_EMTlow_diffexp_notEMT.csv", quote=FALSE, sep="\t",row.names=FALSE)

# output expression values in low_EMT samples
write.table(data.frame(rownames(LIHC.lowEMT),LIHC.lowEMT),"Low_EMT_samples.csv", quote=FALSE, sep="\t",row.names=FALSE)
# output expression values in high_EMT samples
write.table(data.frame(rownames(LIHC.highEMT),LIHC.highEMT),"High_EMT_samples.csv", quote=FALSE, sep="\t",row.names=FALSE)

######
```
